# Chronic Interferon Stimulated Gene Transcription Promotes Oncogene Induced Breast Cancer

**DOI:** 10.1101/2023.10.16.562529

**Authors:** Hexiao Wang, Claudia Canasto-Chibuque, Jun Hyun Kim, Marcel Hohl, Christina Leslie, Jorge S Reis-Filho, John HJ Petrini

## Abstract

The Mre11 complex (comprising Mre11, Rad50, Nbs1) is integral to the maintenance of genome stability. We previously showed that a hypomorphic *Mre11* mutant mouse strain (*Mre11^ATLD1/ATLD1^*) was highly susceptible to oncogene induced breast cancer. Here we used a mammary organoid system to examine which Mre11 dependent responses are tumor suppressive. We found that *Mre11^ATLD1/ATLD1^* organoids exhibited an elevated interferon stimulated gene (ISG) signature and sustained changes in chromatin accessibility. This *Mre11^ATLD1/ATLD1^* phenotype depended on DNA binding of a nuclear innate immune sensor, IFI205. Ablation of *Ifi205* in *Mre11^ATLD1/ATLD1^* organoids restored baseline and oncogene-induced chromatin accessibility patterns to those observed in *WT*. Implantation of *Mre11^ATLD1/ATLD1^* organoids and activation of oncogene led to aggressive metastatic breast cancer. This outcome was reversed in implanted *Ifi205^-/-^ Mre11^ATLD1/ATLD1^* organoids. These data reveal a connection between innate immune signaling and tumor suppression in mammary epithelium. Given the abundance of aberrant DNA structures that arise in the context of genome instability syndromes, the data further suggest that cancer predisposition in those contexts may be partially attributable to tonic innate immune transcriptional programs.

## Introduction

The Mre11 complex controls the DNA damage response (DDR) by governing double-strand break (DSB) repair as well as the activation of the ataxia-telangiectasia mutated (ATM) transducing kinase. ATM activation promotes cell-cycle checkpoint induction, influences DNA repair and induces apoptosis or senescence in particular cellular contexts^1^. Null mutations of *Mre11, Rad50* and *Nbs1* are lethal at the cellular and organismal level^2-4^. Hence, conditional or hypomorphic alleles of Mre11 complex components have been employed to study its role in governing the DDR and in tumor suppression.

We previously queried an array of mice harboring mutations affecting various facets of the DDR network to understand its role in the response to oncogene activation. In *WT* mice, several indices of DDR activation were detected in mammary hyperplasias arising three weeks following induction of *neuT* oncogene, including γH2AX and 53BP1 foci^5^. In *Mre11^ATLD1/ATLD1^* mice, which harbor a hypomorphic *Mre11* mutation inherited in the human ataxia-telangiectasia like disorder (A-TLD)^6^, these DDR outcomes were abolished. This phenotype correlated with much more extensive hyperplasias three weeks post oncogene activation. Whereas progression to tumors in *WT* mice was rare (5%), the florid hyperplasias in *Mre11^ATLD1/ATLD1^* mice presaged the frequent (40%) onset of highly aggressive basal like tumors. The extent of hyperplasia and formation of DDR foci were unaffected by mutations that disrupted apoptosis (*Trp53^515C/515C^*, *Chk2^-/-^*) and DNA repair (*53BP1^-/-^*), and tumor latency was indistinguishable from *WT* in those genetic contexts^5^.

The previous study also revealed an unrecognized role for the Mre11 complex in oncogene-induced chromatin modifications. *neuT* expressing *WT* mammary tissues exhibited deposition of heterochromatic markers such as macroH2A and phospho histone H3 (S10) accompanied by an arrest in the G2 phase of the cell cycle. Unlike the formation of DDR foci, these events required both the Mre11 complex and p53^5^. It is not clear which of the downstream functions of Mre11 and p53 are tumor suppressive in this context.

Interferon inducible gene 205 (*Ifi205*) belongs to the HIN-200 family, and is a regulator of type I interferon (IFN) production^7,8^. HIN-200 family members are characterized by at least one HIN-200 domain (hematopoietic interferon-inducible nuclear proteins with a 200-amino-acid repeat), which binds double-stranded DNA via an oligonucleotide/oligosaccharide binding (OB) fold^9,10^. HIN-200 family members also contain an N terminal pyrin domain, which mediates assembly into innate immune effector complexes^8,11^. Most HIN-200 proteins possess a nuclear localization signal and localize to the nucleus^12^. While the HIN-200 family is believed to be involved in the modulation of type I IFN and inflammasome pathways, the precise roles of most of the individual genes have not been elucidated^7,8^.

The cancer susceptibility in *Mre11^ATLD1/ATLD1^* mice could be attributable to an Mre11 complex dependent pathway which is activated upon oncogenic stress. An alternative possibility is that Mre11 complex hypomorphism somehow creates a permissive state for oncogene induced carcinogenesis. We employed an organoid system comprising primary mammary epithelial cells (MECs) that harbor an inducible oncogene, *neuT*, to examine these possibilities. The organoid system enables *ex vivo* manipulation and reimplantation to query the effects of *ex vivo* genetic manipulations on tumor development.

In the basal state (*i.e.,* prior to oncogene activation)*, Mre11^ATLD1/ATLD1^* organoids exhibited a tonic interferon stimulated gene (ISG) signature. This coincided with changes in the accessibility of numerous chromatin loci. The ISG transcriptional signature and the changes in chromatin accessibility at baseline were dependent on IFI205 DNA binding. As observed in the autochthonous mammary analyses noted above^5^, oncogene induction in implanted *Mre11^ATLD1/ATLD1^* organoids led to aggressive basal like breast tumors. Ablation of *Ifi205* in *Mre11^ATLD1/ATLD1^*organoids reverted chromatin accessibility to that of *WT* and restored the oncogene-induced heterochromatic changes observed in *WT* upon oncogene induction. This correlated with a sharp reduction in tumor frequency and increased latency in implanted *Ifi205^-/-^ Mre11^ATLD1/ATLD1^* organoids relative to *Mre11^ATLD1/ATLD1^* organoids. This study is consistent with the interpretation that oncogene induced chromatin changes previously observed in *WT* MECs are tumor suppressive, and suggests a previously unrecognized link between ISG transcription and tumor suppression.

## Results

We generated *WT* and *Mre11^ATLD1/ATLD1^* mice that contain an inducible *neuT* gene. *CAG-rtTA* transgenic mice^13^, in which reverse tetracycline-controlled trans activator (*rtTA*) is driven by a tissue non-specific *CAG* promoter, were crossed with *TetO-neuT* transgenic mice^14^, allowing for doxycycline-inducible expression of the oncogene *neuT* and a linked luciferase reporter (Fig.1A). Primary mammary epithelial cells (MECs) were harvested for the establishment of primary mammary organoids adapting previously described methods for intestinal and prostate epithelia^15,16^ (Fig.1B). This system allows us to control oncogene activation in the organoids *ex vivo* and achieve mammary tissue specific oncogene activation *in vivo* once the organoids are implanted into recipient mice.

**Figure 1.**
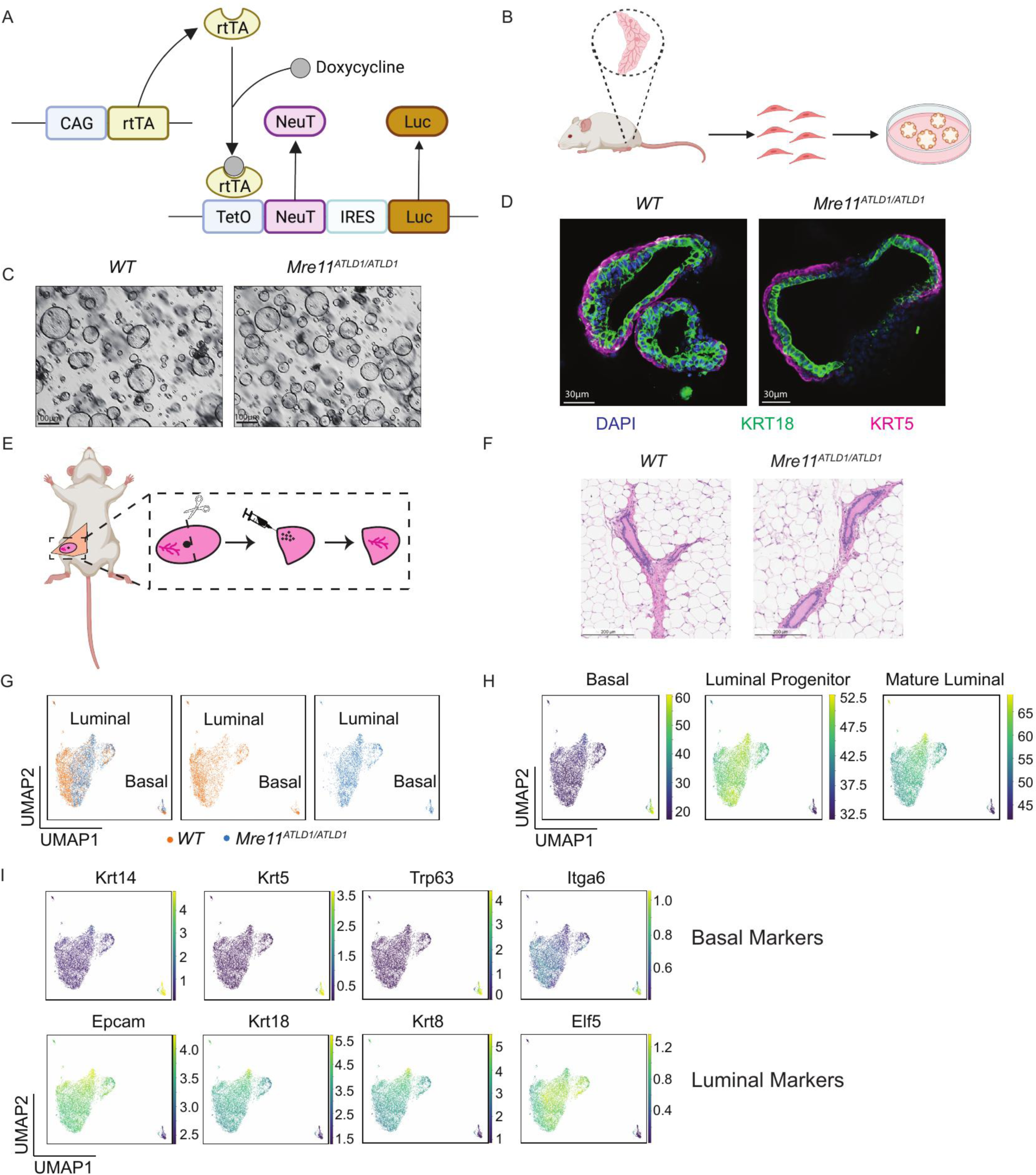
Mammary epithelia derived from W7 and *Mre11^ATLD1/ATLD1^* mice develop into normal organoids. (A) Schematic representation of doxycycline inducible *neuT* and luciferase reporter transgene. (B) Experimental strategy for primary mammary epithelial cell isolation and mammary organoid culture. (C) Bright field images of *WT* and *Mre11^ATLD1/ATLD1^* mammary organoids. (D) Immunofluorescence (IF) staining of lineage markers (KRT18 and KRT5) in *WT* and *Mre11^ATLD1/ATLD1^*mammary organoids. (E) Strategy of mammary fat pad clearance and organoid implantation. (F) Hematoxylin and eosin (H&E) staining of mammary tissues six weeks after *WT* and *Mre11^ATLD1/ATLD1^* organoid implantation. (G) Uniform Manifold Approximation and Projection (UMAP) plot color-coded by genotype from single cell RNA-Seq in *WT* and *Mre11^ATLD1/ATLD1^* mammary organoids. (H) UMAP plot color-coded by cell lineage signature score from single cell RNA-Seq in *WT* and *Mre11^ATLD1/ATLD11^* mammary organoids. (I) UMAP plot color-coded by expression level of cell lineage markers as indicated from single cell RNA-Seq in *WT* and *Mre11^ATLD1/ATLD1^* mammary organoids.

### Organoids derived from WT and Mre11^ATLD1/ATLD1^ mice develop normal mammary glands

The conditions established for mammary organoids were compatible with normal mammary gland development. Organoids from both *WT* and *Mre11^ATLD1/ATLD1^* MECs formed hollow spheres, reminiscent of mammary gland architecture *in vivo* (Fig.1C). Immunofluorescence (IF) staining showed that both *WT* and *Mre11^ATLD1/ATLD1^* organoids contained KRT5 positive basal cells and KRT18 positive luminal cells (Fig.1D). *WT* and *Mre11^ATLD1/ATLD1^*organoids formed histologically normal mammary glands *in vivo* after implantation into cleared mammary fat pads of NOD-SCID mice (Fig.1E-F).

The composition of the mammary organoids was further assessed by single cell RNA-Seq. Transcriptomes from 7393 cells (3850 *WT*, 3543 *Mre11^ATLD1/ATLD1^*) were analyzed and visualized by Uniform Manifold Approximation and Projection (UMAP). The gene expression profiles revealed that luminal clusters and basal clusters were evident in *WT* and *Mre11^ATLD1/ATLD1^* organoids (Fig.1G). Those clusters were verified using common lineage markers (*Krt14, Krt5, Trp63, Itga6* for basal cells and *Epcam, Krt18, Krt8, Elf5* for luminal cells) or lineage signature transcriptional score generated from normal mammary tissues^17^ (Fig.1H-I). These data demonstrate that *WT* and *Mre11^ATLD1/ATLD1^* organoids are indistinguishable in the competency for mammary gland development.

### WT and Mre11^ATLD1/ATLD1^ organoids show different oncogenic responses after neuT activation

Having verified that *WT* and *Mre11^ATLD1/ATLD1^* organoids exhibited the same cellular composition of mammary glands *in vivo*, we assessed their responses to oncogene activation. Doxycycline was added to organoid cultures to induce *neuT* expression. Western blot confirmed that *neuT* was induced to similar levels in *WT* and *Mre11^ATLD1/ATLD1^* organoids (Fig.S1). As observed *in vivo*^5^, oncogene activation induced DDR and heterochromatic markers in *WT* but not *Mre11^ATLD1/ATLD1^* organoids. Western blotting was performed at day 0, day 14 and day 28 after adding doxycycline. γH2AX was induced at day 28 after *neuT* activation in *WT* organoids, and heterochromatic markers Hp1-γ and macroH2A2, also increased after oncogene activation (Fig.2A). In contrast, *Mre11^ATLD1/ATLD1^* organoids did not show induction of γH2AX after oncogene activation and the induction of Hp1-γ and macroH2A2 was also greatly diminished, consistent with prior findings that induction of DDR and chromatin markers are dependent on the Mre11 complex (Fig.2A). Hence, the mammary organoid system adapted here recapitulates previously observed responses to oncogene *in vivo*^5^.

**Figure 2.**
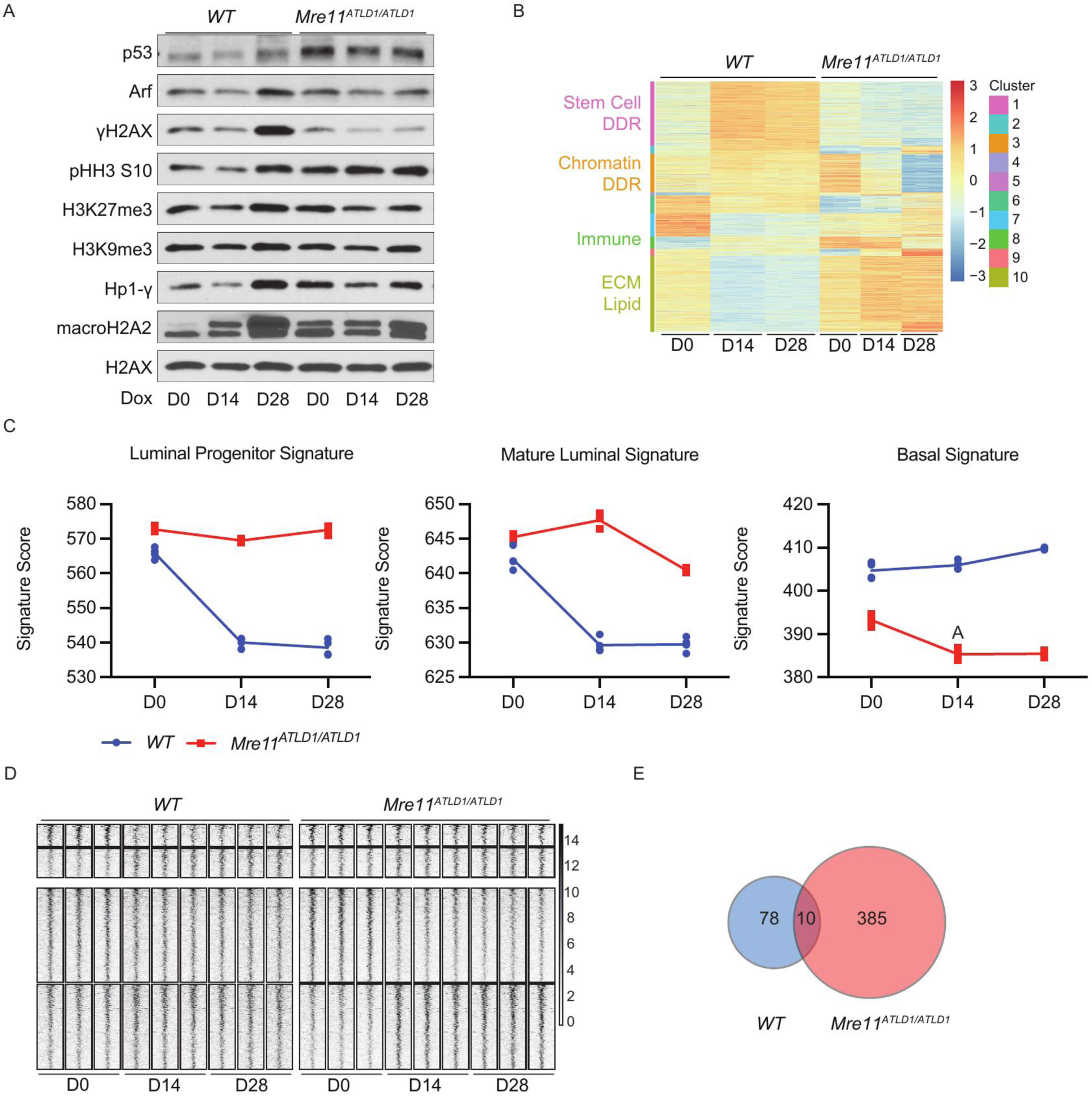
*WT* and *Mre11^ATLD1/ATLD1^* organoids show different oncogenic responses after *neuT* activation. (A) Western blot of whole cell extracts taken from *WT* and *Mre11^ATLD1/ATLD1^* organoids at indicated timepoints after adding 1μg/ml doxycycline. (B) Heatmap of 5644 genes exhibiting different patterns of changes after oncogene activation between *WT* and *Mre11^ATLD1/ATLD1^* organoids from bulk RNA-Seq analysis (padj ≤ 0.01, |FC| ≥ 1.5). Labels on the left summarize the corresponding cluster based on pathway and biological function analysis. Stem Cell and DNA Damage Response (DDR) for cluster 1, Chromatin and DDR for cluster 3, Immune for cluster 8, Extracellular Matrix (ECM) and Lipid for cluster 10. (C) Luminal progenitor signature score, mature luminal signature score and basal signature score calculated from bulk RNA-Seq in *WT* and *Mre11^ATLD1/ATLD1^* organoids at indicated timepoints after adding 1 pg/ml doxycycline. (D) Heatmap of chromatin regions (rows) showing different accessibility at indicated timepoints after adding 1μg/ml doxycycline in *WT* and *Mre11^ATLD1/ATLD1^* organoids from ATAC-Seq analysis. Top, chromatin regions showing different accessibility after oncogene activation in *WT* organoids (left), status of the same regions at the same timepoints in *Mre11^ATLD1/ATLD1,^* organoids were plotted on the right as comparison (88 peaks, padj ≤ 0.05, |FC| ≥ 1.5). Bottom, chromatin regions showing different accessibility after oncogene activation in *Mre11^ATLD1/ATLD1^* (right), status of the same regions at the same timepoints in *WT* organoids were plotted on the left as comparison (395 peaks, padj ≤ 0.05, |FC| ≥ 1.5). (E) Venn diagram of chromatin regions showing oncogene-induced changes in accessibility in *WT* and *Mre11^ATLD1/ATLD1,^* organoids from (D).

To examine the potential mechanistic bases for Mre11 complex dependent tumor suppression in mammary epithelium, the responses of *WT* and *Mre11^ATLD1/ATLD1^* organoids to oncogene activation were assessed. RNA was prepared from *WT* and *Mre11^ATLD1/ATLD1^* organoids at day 0, day 14, day 28 after doxycycline induction, followed by bulk RNA-Seq. 5,644 genes exhibited different changes in expression after oncogene activation between *WT* and *Mre11^ATLD1/ATLD1^* (padj ≤ 0.01, |FC| ≥ 1.5). Among them, approximately 30% (1539/5644) were already differentially expressed prior to oncogene activation. Clustering and pathway analysis of genes with different expression patterns revealed signatures related to Stem Cell, DDR, Chromatin, Immune Response, Extracellular Matrix (ECM) and Lipid Metabolism (Fig.2B). While *WT* and *Mre11^ATLD1/ATLD1^*organoids displayed similar lineage signature scores at baseline, both luminal progenitor signature score and mature luminal signature score decreased significantly in response to oncogene activation in *WT* organoids, whereas a reduction of basal signature score was observed in *Mre11^ATLD1/ATLD1^* organoids, indicating different oncogene-induced trajectories may exist in *WT* and *Mre11^ATLD1/ATLD1^*organoids (Fig.2C). This is consistent with our previous finding that *neuT* induced *Mre11^ATLD1/ATLD1^* tumors are basal-like breast cancer^5^, which have been reported to originate from the luminal progenitors^18,19^.

Given the difference in oncogene-induced chromatin modifications between *WT* and *Mre11^ATLD1/ATLD1^*organoids, the effect of oncogene activation on chromatin accessibility was assessed. ATAC-Seq was performed using *WT* and *Mre11^ATLD1/ATLD1^* organoids at the same timepoints as above. Chromatin accessibility analysis revealed substantial differences in response to oncogene activation between these two genotypes. In *WT* organoids, 0.2% of the total chromatin regions (64/22,950) became more open and 0.1% regions (24/22,950) became more closed after oncogene activation (Fig.2D top). In contrast, *Mre11^ATLD1/ATLD1^* organoids exhibited greater changes in chromatin accessibility, with 0.8% chromatin regions (187/22,950) becoming more accessible and 0.9% regions (208/22,950) becoming less accessible after oncogene activation (Fig.2D bottom). However, only 10 regions showed similar changes in accessibility in both *WT* and *Mre11^ATLD1/ATLD1^* organoids, underscoring divergent oncogene associated chromatin responses in those genotypes (Fig.2E).

### Mre11^ATLD1/ATLD1^ organoids exhibit tonic ISG transcriptional signature and altered chromatin status at basal state

The differential responses and increased tumorigenesis in *neuT* expressing *Mre11^ATLD1/ATLD1^* MECs can be explained by two non-exclusive possibilities. First, the Mre11 complex may control tumor-suppressive pathways that are induced by oncogenic stress. Alternatively, Mre11 complex hypomorphism may create a state of permissiveness for oncogene-driven carcinogenesis. We reasoned that comparison of *WT* and *Mre11^ATLD1/ATLD1^* organoids at basal state would address the latter possibility.

Bulk RNA-Seq was performed in *WT* and *Mre11^ATLD1/ATLD1^* organoids, followed by canonical pathway analysis and Gene Set Enrichment Analysis (GSEA). Prior to oncogene activation, an interferon stimulated gene (hereafter ISG) transcriptional program was the most significant difference between *WT* and *Mre11^ATLD1/ATLD1^* organoids (Fig.3A-B). The ISG pathway observed in *Mre11^ATLD1/ATLD1^*organoids reflected upregulation of multiple ISGs (*e.g., Nos2, Ccl20, Ifi205, Ccl5, Ifit1(Isg56), Isg15, Irf7, Stat1*) (Fig.3C), leading to a stronger ISG transcriptional signature. Single-cell RNA-Seq also showed a uniform increase in the ISG signature score in *Mre11^ATLD1/ATLD1^* organoids, indicating that the enhanced signature seen in the bulk RNA-Seq does not originate from a sub-population of cells (Fig.3D, Fig.S2).

**Figure 3.**
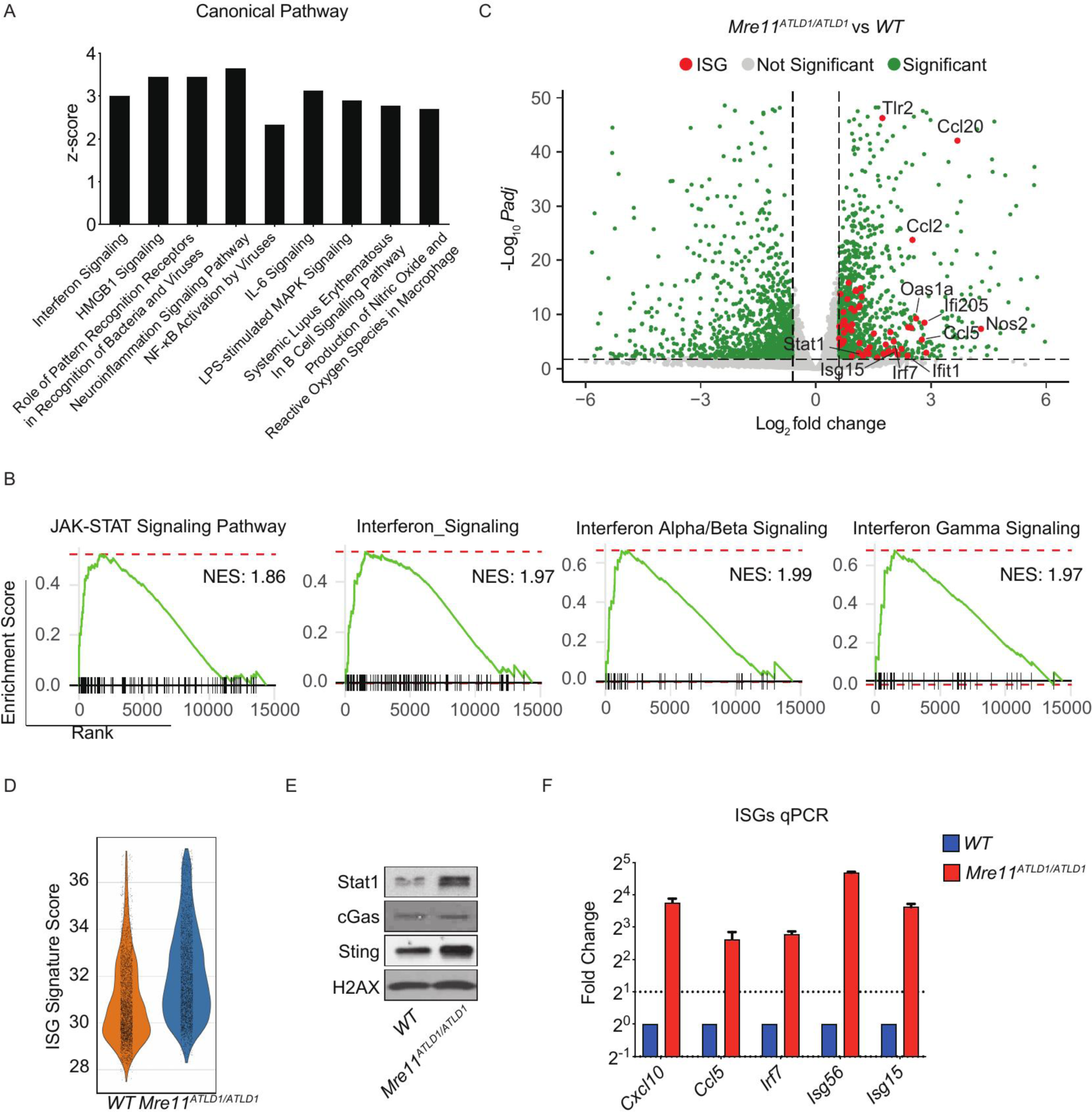
*Mre11^ATLD1/ATLD1^* organoids show stronger ISG signature at basal state. (A) Canonical pathway analysis performed by Ingenuity Pathway Analysis (IPA) using differentially expressed genes (DEGs, Padj ≤ 0.05, |FC| ≥ 1.5) from comparison between *Mre11^ATLD1/ATLD1^* and *WT* organoids at baseline in RNA-Seq (Padj ≤ 0.05). Z-score was calculated by comparing *Mre11^ATLD1/ATLD1^* vs *WT.* The positive z-score predicts activation state of the pathway. An absolute z-score a 2 is considered significant. (B) Gene Set Enrichment Analysis (GSEA) comparison between *Mre11^ATLD1/ATLD1^* and *WT* organoids at baseline in RNA-Seq. (C) Volcano plot from comparison between *Mre11^ATLD1/ATLD1^* and *WT* organoids at baseline in RNA-Seq. Green, significant (Padj ≤ 0.05, |FC| ≥ 1.5); grey, not significant; red, significant ISGs. (D) Violin plot of ISG signature score from single cell RNA-Seq in *WT* and *Mre11^ATLD1/ATLD1^* organoids. (E) Western blot of whole cell extracts taken from *WT* and *Mre11^ATLD1/ATLD1^* organoids. (F) Expression levels of ISGs measured by qPCR in *WT* and *Mre11^ATLD1/ATLD1^* organoids.

Orthogonal validation of the RNA-Seq analysis was obtained from Western blotting and qPCR. Western blot analysis showed increased protein levels of Stat1 and Sting in *Mre11^ATLD1/ATLD1^* organoids compared with *WT* (Fig.3E). qPCR also revealed up- regulation of common downstream ISGs (*e.g.*, *Cxcl10*, *Ccl5, Irf7, Isg56, Isg15*) in *Mre11^ATLD1/ATLD1^* organoids (Fig.3F). These data confirm the presence of a stronger ISG signature in *Mre11^ATLD1/ATLD1^* organoids compared with *WT* organoids at baseline prior to oncogene activation.

Chronic ISG transcription has been shown to trigger widespread changes in chromatin accessibility^20^. We therefore investigated the potential effect of ISG transcriptional program on chromatin status in the organoids. Bulk RNA-Seq analysis revealed that 36 chromatin modifiers and remodelers (*e.g., Jade2, Kat2b, Foxa1* and *Hmgn5*) were differentially expressed in *Mre11^ATLD1/ATLD1^* compared with *WT* organoids (Fig.4A, padj ≤ 0.01, |FC| ≥ 1.5). Notably, 18 of these chromatin regulators are known to be regulated by interferon (IFN) signaling pathway^21-38^ (asterisk in Fig.4A, Table S1), which likely impart chromatin changes due to the tonic ISG transcriptional program in *Mre11^ATLD1/ATLD1^*organoids. ATAC-Seq analysis was performed in *WT* and *Mre11^ATLD1/ATLD1^* organoids at baseline to assess difference in chromatin accessibility. The accessibility of 2.0% of chromatin regions (456/22,950) were different between the two genotypes; 0.5% of the regions (118/22,950) more open and 1.5% (338/22,950) more closed in *Mre11^ATLD1/ATLD1^* organoids (Fig.4B, padj ≤ 0.05, |FC| ≥ 1.5).

**Figure 4.**
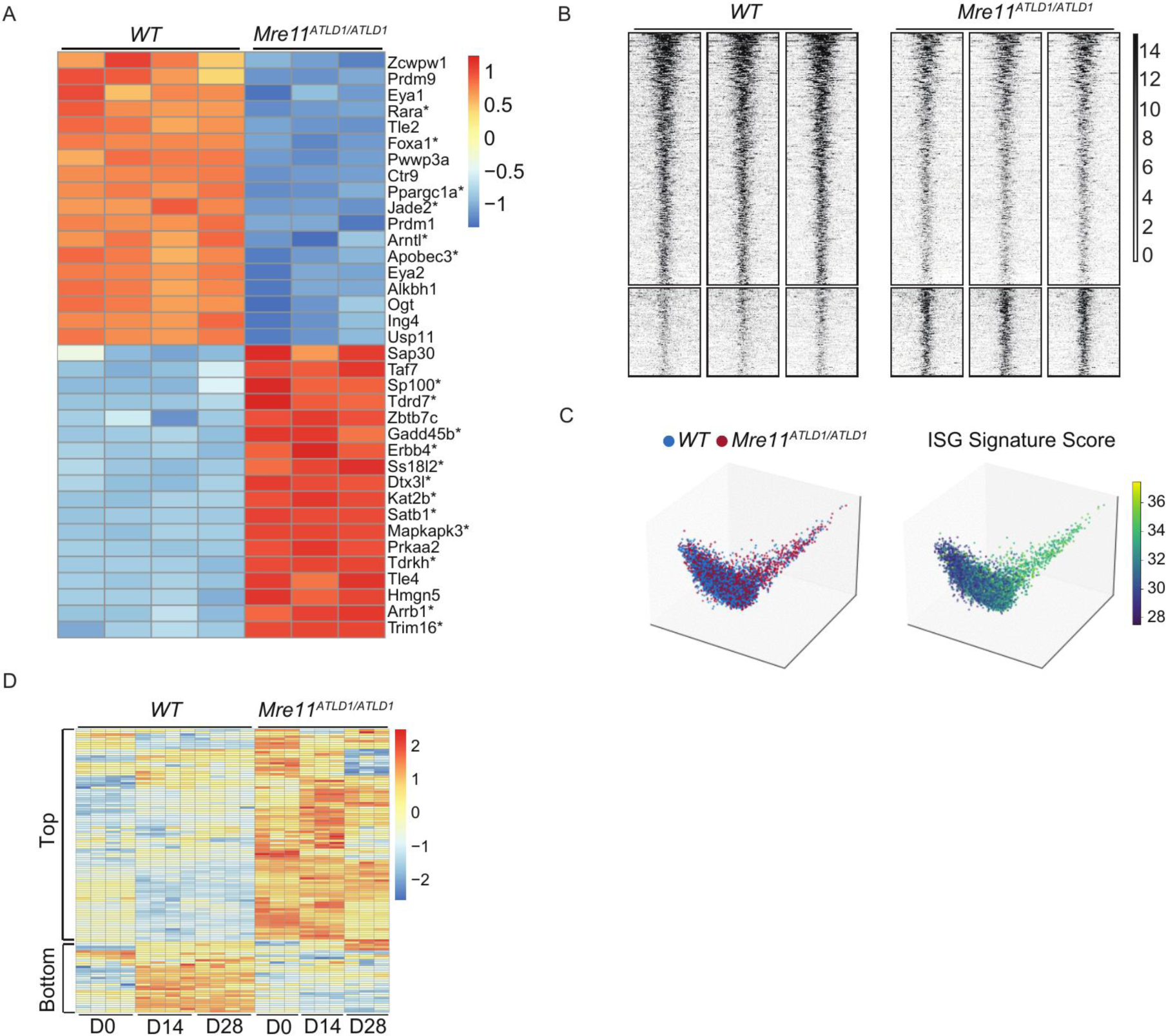
*Mre11^ATLD1/ATLD1^* dependent ISG signature correlates different chromatin status at basal state. (A) Heatmap of differentially expressed chromatin modifiers and remodelers from RNA-Seq in *WT* and *Mre11^ATLDVATLD1^* organoids (padj ≤ 0.01, |FC| a 1.5). (B) Heatmap of differential chromatin accessibility regions (rows) between *WT* and *Mre11^ATLD1/ATLD1^* organoids from ATAC-Seq (Padj ≤ 0.05, |FC| ≥ 1.5, 118 regions more accessible and 338 regions less accessible in *Mre11^ATLD1/ATLD1^).* (C) Potential of Heat-diffusion for Affinity-based Trajectory Embedding (PHATE) 3D plot using chromatin modifiers and remodelers color-coded by genotype (left) and ISG signature score (right) from single cell RNA-Seq in *WT* and *Mre11^ATLD1/ATLD1^* organoids. (D) Heatmap of a list of ISG (GO: cellular response to interferon-alpha, beta and gamma) expression levels from RNA-Seq in *WT* and *Mre11^ATLD1/ATLD1^*organoids at day 0, day 14, day 28 after oncogene activation. Top group represents up-regulated ISGs in *Mre11^ATLD1/ATLD1^* organoids at baseline which were remained at higher levels after oncogene activation. Bottom group represents induced ISGs in *WT* organoids after oncogene activation whose induction was abolished in *Mre11^ATLD1/ATLD1^* organoids.

Single-cell RNA-Seq suggested that the ISG signature observed correlated with chromatin status in *Mre11^ATLD1/ATLD1^* organoids. To visualize chromatin status and ISG signature simultaneously at single-cell level, expression levels of 113 chromatin modifiers and remodelers examined in a previous study^39^ were used as a proxy of chromatin status. Potential of Heat-diffusion for Affinity-based Trajectory Embedding (PHATE) analysis was performed, which was then color-coded by ISG signature score. The resulting 3D PHATE plot revealed a distinct group of cells with differential expression of chromatin modifiers and remodelers (Fig.4C left) and higher ISG signature scores (Fig.4C right), suggesting a correlation between chromatin status and ISG signatures. These findings demonstrate that the ISG transcriptional program found in *Mre11^ATLD1/ATLD1^* organoids at basal level correlates with changes in chromatin status.

Oncogene activation causes DNA damage that is likely linked to DNA replication stress^40-42^. In turn, DNA damage can lead to ISG induction^43,44^. To evaluate the status of ISG expression after oncogene activation, we analyzed a list of ISGs from Gene Ontology (GO, cellular response to interferon-alpha, beta and gamma, Table S2) using bulk RNA-Seq data in *WT* and *Mre11^ATLD1/ATLD1^* organoids after doxycycline addition. Unsupervised clustering of these genes revealed two major groups. A unique subset of ISGs were induced in *WT* organoids by oncogene activation (Fig.4D bottom group). In contrast, the ISG signature observed in *Mre11^ATLD1/ATLD1^*organoids at baseline remained at higher levels and was largely unchanged upon oncogene activation (Fig.4D top group). That the subset of genes induced in *WT* organoids is distinct from the subset elevated in *Mre11^ATLD1/ATLD1^*may indicate that distinct upstream factors underlie the respective ISG signatures. These data indicate that the *Mre11^ATLD1/ATLD1^* genotype, as opposed to oncogene activation was the primary determinant of the ISG signature in those organoids.

### Ifi205 induces ISG signature and mediates chromatin changes in Mre11^ATLD1/ATLD1^ organoids

The ISG signature observed in *Mre11^ATLD1/ATLD1^* organoids was not attributable to cGas-Sting pathway. As previously observed in *Mre11^ATLD1/ATLD1^* MEFs^45^, spontaneous chromosome aberrations were elevated in *Mre11^ATLD1/ATLD1^* organoids compared with *WT* (Fig.5A-B, p=0.0485, aberrations: *WT* 0/21, *Mre11^ATLD1/ATLD1^* 5/22). However, no significant increase of cytoplasmic dsDNA or cGAMP was detected in *Mre11^ATLD1/ATLD1^*organoids compared with *WT* (Fig.S3), indicating that cGas-Sting cytosolic DNA sensing is not the major pathway responsible for the ISG signature in *Mre11^ATLD1/ATLD1^*organoids.

**Figure 5.**
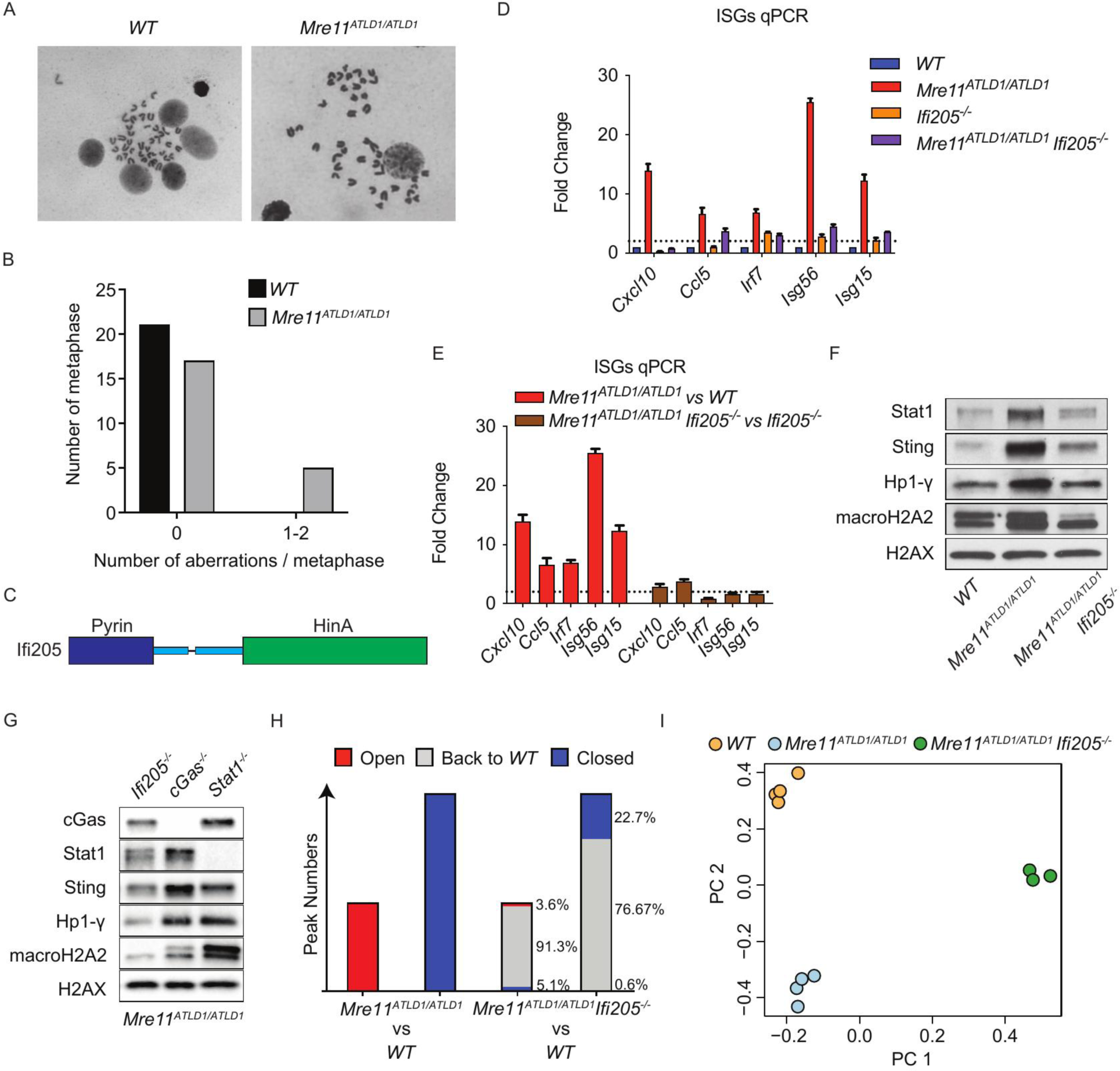
IFI205 induces ISG signature and mediates chromatin changes in *Mre11^ATLD1/ATLD1^* organoids. (A) Example images of metaphase spreads in *WT* and *Mre11^ATLD1/ATLD1^* organoids. (B) Quantification of metaphase spreads in A (Fisher’s exact test, p = 0.0485; aberrations: M/T 0/21, *Mre11^ATLD1/ATLD1^* 5/22). (C) Schematic representation of IFI205 protein. (D) Expression levels of ISGs measured by qPCR in *WT, Mre11^ATLD1/ATLD1^, lfi205^-/-^and lfi205^-/-^ Mre11^ATLD1/ATLD1^* organoids. Fold changes were calculated as *Mro11^ATLD1/ATLD1^ vs WT, lfi205^-/-^* vs *WT,* and *lfi205^-/-^ Mre11^ATLD1/ATLD1^* vs *WT.* (E) Same as (D). Fold changes were calculated as *Mre11^ATLD1/ATLD1^* WT, and *lfi205^-/-^ Mre11^ATLD1/ATLD1^* vs *lfi205v* (F) Western blot of whole cell extracts taken from *WT,* Mre11^ATLD1/ATLD1^ and *lfi205^-/-^ Mre11^ATLDVATLD1^ organoids.* (G) Western blot of whole cell extracts taken from *lfi205^-/-^ Mre11^ATLD1/ATLD1^, cGas^-/-^ Mre11^ATLD1/ATLD1^* and *Stat1^-/-^ Mre11^ATLD1/ATLD1^* organoids. (H) Quantification of *lfi205^-/-^ Mre11^ATLD1/ATLD1^* W*T* organoids comparison from ATAC-Seq, using differential chromatin accessibility regions between *Mre11^ATLDVATLD1^* and *WT* organoids (Open, padj ≤ 0.05 and FC ≥ 1.5; Closed, padj ≤ 0.05 and FC ≤ -1.5). (I) Principal Component Analysis (PCA) plot from ATAC-Seq in *WT, Mre11^ATLDlaTLD1^* and *lfi205^-/-^ Mre11^ATLD1/ATLD1^* organoids (Open, padj ≤ 0.05 and FC ≥ 1.5; Closed, padj ≤ 0.05 and FC ≤ -1.5).

Given that cytoplasmic DNA does not appear to underlie the observed ISG signature, we hypothesized that a nuclear sensor may engage aberrant DNA structures and induce the observed tonic ISG signature. We found that the transcript encoding the nuclear DNA sensor IFI205 was up-regulated in *Mre11^ATLD1/ATLD1^* organoids (Fig.3C, FC = 7.11, padj = 3.3676x10^-9^). This gene product was also recently found to be enriched at stressed DNA replication forks^46^.

IFI205 binds dsDNA through its HIN domain and activates IFN signaling^12,47^ (Fig.5C). To determine if the ISG signature and subsequent chromatin changes were dependent on Ifi205, we used CRISPR-Cas9 to inactivate the *Ifi205* gene in *WT* and *Mre11^ATLD1/ATLD1^*organoids. qPCR showed that while only *Cxcl10* level decreased in *Ifi205^-/-^* organoids compared to *WT* organoids (FC = 0.27), all the ISGs that were elevated in *Mre11^ATLD1/ATLD1^* (*Cxcl10*, *Ccl5, Irf7, Isg56, Isg15*) were reduced after knocking-out *Ifi205* in *Mre11^ATLD1/ATLD1^* organoids (Fig.5D-E; FC, *Ifi205^-/-^ Mre11^ATLD1/ATLD1^* vs *Mre11^ATLD1/ATLD1^*, *Cxcl10* = 2.91, *Ccl5* = 3.78*, Irf7* = 0.89*, Isg56* = 1.59*, Isg15* =1.66). Western blot showed that both IFN related proteins (Stat1 and Sting) and chromatin markers (Hp1-γ and macroH2A2) decreased in *Mre11^ATLD1/ATLD1^ Ifi205^-/-^* organoids compared with *Mre11^ATLD1/ATLD1^* organoids (Fig.5F). In contrast, knocking-out *cGas* or *Stat1* in *Mre11^ATLD1/ATLD1^* organoids failed to inhibit Sting, Hp1-γ and macroH2A2 protein levels to that observed in *Ifi205^-/-^ Mre11^ATLD1/ATLD1^* organoids (Fig.5G).

Ablation of *Ifi205* also largely reverted *Mre11^ATLD1/ATLD1^* chromatin accessibility to *WT*. ATAC-Seq analysis showed that 91.3% of the previously more open chromatin regions and 76.7% of the previously more closed chromatin regions in *Mre11^ATLD1/ATLD1^*organoids reverted to *WT* status in *Ifi205^-/-^ Mre11^ATLD1/ATLD1^* organoids (Fig.5H, *Ifi205^-/-^ Mre11^ATLD1/ATLD1^* vs *WT,* padj > 0.05 or |FC| < 1.5). Principle Component Analysis (PCA) of ATAC-Seq also showed that, on PC2, *Ifi205^-/-^ Mre11^ATLD1/ATLD1^* exhibited a trend of shifting back to *WT* chromatin (Fig.5I).

In addition to the normalization of baseline ISG signature and chromatin accessibility, IFI205 deficiency in *Mre11^ATLD1/ATLD1^* organoids also restored the response to oncogene activation. Western blot was performed using samples collected at day 0, day 14 and day 28 after adding doxycycline from *WT, Mre11^ATLD1/ATLD1^* and *Ifi205^-/-^ Mre11^ATLD1/ATLD1^* organoids. Induction of macroH2A2 and Hp1-γ could be detected in *Ifi205^-/-^ Mre11^ATLD1/ATLD1^* organoids after oncogene activation similar to that in *WT* (Fig.6A). To assess alterations in chromatin accessibility after oncogene activation, ATAC-Seq was performed at same timepoints in *WT, Mre11^ATLD1/ATLD1^* and *Ifi205^-/-^ Mre11^ATLD1/ATLD1^* organoids. PCA derived from oncogene-induced chromatin regions identified in *WT* and *Mre11^ATLD1/ATLD1^* organoids, revealed more similar trends between *Ifi205^-/-^ Mre11^ATLD1/ATLD1^* and *WT* organoids than that between *Ifi205^-/-^ Mre11^ATLD1/ATLD1^* and *Mre11^ATLD1/ATLD1^*organoids, suggesting *Ifi205^-/-^ Mre11^ATLD1/ATLD1^* largely restores oncogene-induced chromatin changes observed in *WT* (Fig.6B). These data demonstrate that IFI205 underlies the ISG signature and subsequent chromatin changes observed in *Mre11^ATLD1/ATLD1^* organoids. Ablation of *Ifi205* in *Mre11^ATLD1/ATLD1^* organoids not only reverts chromatin accessibility to *WT* at baseline, but also recovers the induction of chromatin changes after oncogene activation.

**Figure 6.**
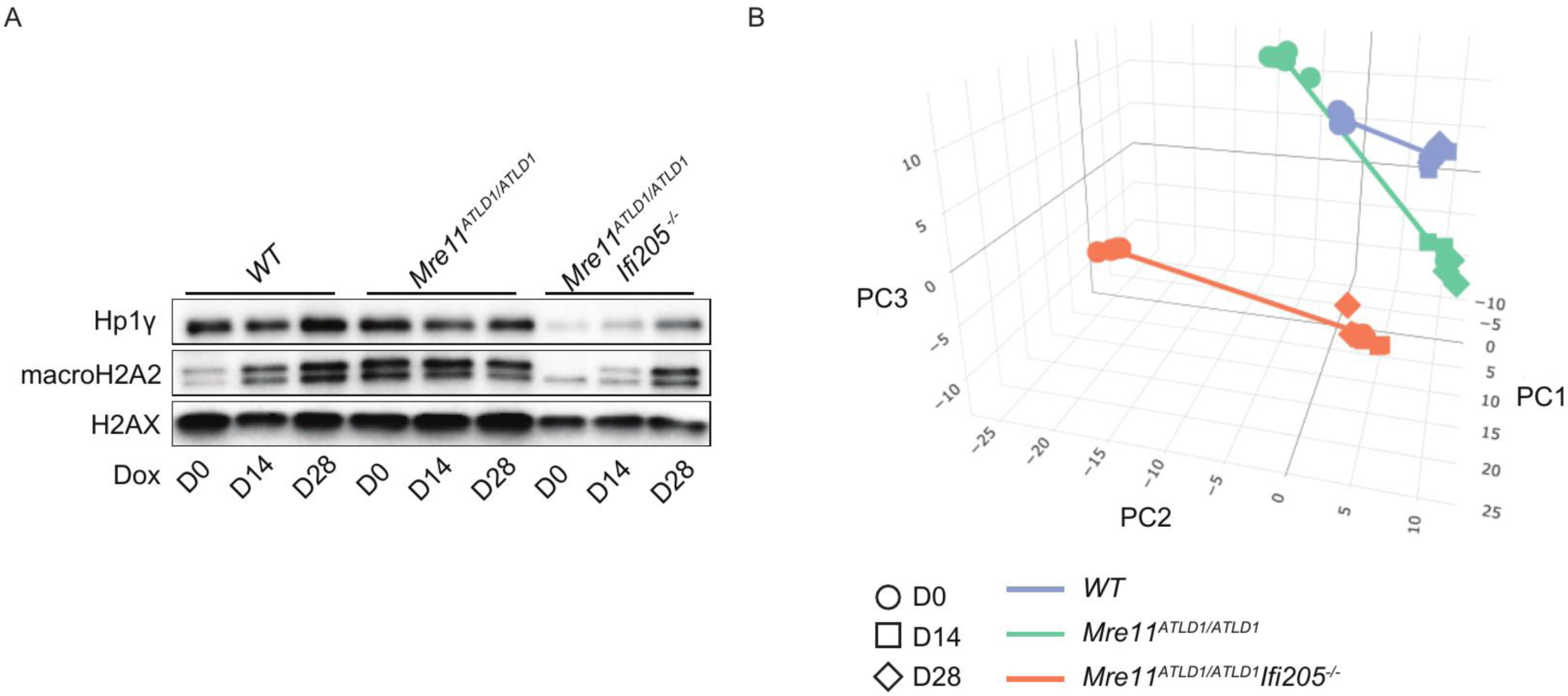
Inactivation of *lfi205* in *Mre11^ATLD1/ATLD1^* organoids restores oncogene-induced chromatin changes. (A) Western blot of whole cell extracts taken from *WT, Mre11^ATLD1/ATLD1,^* and *lfi205^-/-^ Mre11^ATLD1/ATLD1^* organoids at indicated timepoints after adding 1 μg/ml doxycycline. (B) PCA plot from ATAC-Seq in *WT, Mre11^ATLD1/ATLD1^* and *lfi205^-/-^ Mre11^ATLD1/ATLD1^* organoids at indicated timepoints after adding 1μg/ml doxycycline, using regions showed oncogene-induced changes in *WT* and *Mre11^ATLDWTLD1^* organoids.

### Ifi205 triggers ISG signature and chromatin changes through DNA sensing

Based on the outcomes of *Ifi205* deficiency in *Mre11^ATLD1/ATLD1^*MECs, we hypothesized that IFI205 senses aberrant DNA structures (*e.g.*, extrachromosomal DNA or decondensed chromatin) in *Mre11^ATLD1/ATLD1^* organoids and promotes both the ISG transcriptional signature and the attendant chromatin changes. This proposal explicitly predicts that DNA binding by IFI205 underlies the effect of *Ifi205* on the *Mre11^ATLD1/ATLD1^*phenotype. Therefore, we designed a DNA binding deficient *Ifi205* mutant to test this hypothesis. Previous study showed six mutations in AIM2 HIN domain disrupted DNA binding ability^48^, among which, two sites (K248 and K313) were facing towards DNA in the structure^49^ (Fig.7A), and were conserved within the IFI205 HIN domain (R299 and K364) (Fig.7B). Hence, the *Ifi205* mutant (*Ifi205^K364ER299E^*) harboring both K364E and R299E mutations was constructed.

**Figure 7.**
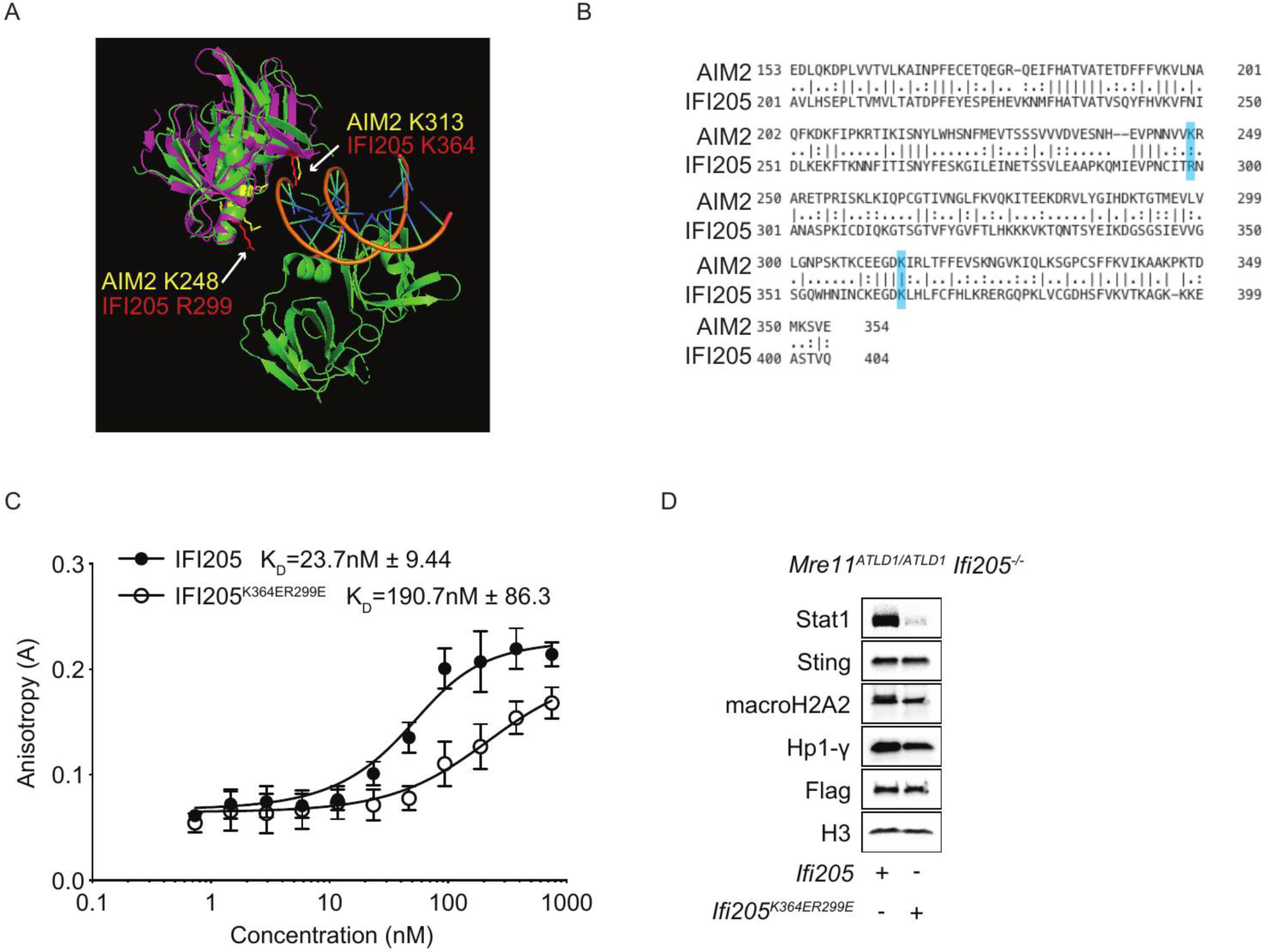
IFI205 triggers ISG signature and chromatin changes through DNA sensing. (A) Structure of AIM2 HIN domain with dsDNA(PDB: 4jbm) overlaid by IFI205 structure predicted by Phyre2. Mutataions modeled later in IF 1205 were colored in red and correspondingly conserved sites in AIM2 were colored in yellow. (B) Sequence alignment between AIM2 and IFI205. Mutatations modeled later in IFI205 were colored in blue, (line, fully conserved; colon, strongly similar; dot, weakly similar) (C) Binding curves from fluorescence anisotropy using 60mer FAM-labeled dsDNA at 50nM and increasing amount of recombinant IFI205 or IFI205^K364ER299E^ protein. (D) Western blot of whole cell extracts taken from *lfi205^-/-^ Mre11^ATLD1/ATLD1^* organoids ectopically expressing either *lfi205* or *lfi205^K364ER299E^*.

DNA binding was assessed using Electrophoretic Mobility Shift Assay (EMSA) and fluorescence anisotropy. Recombinant FLAG-tagged IFI205 and IFI205^K364ER299E^ were purified from *E.coli* (Fig.S4). We measured DNA binding affinity of IFI205 and IFI205^K364ER299E^ by fluorescence anisotropy, using 50nM FAM labeled 60mer dsDNA^50^. The K_D_ of IFI205^K364ER299E^ was 190.7nM±86.3, eight fold higher than that of IFI205 (23.7nM±9.44), indicating a significant decrease of DNA binding affinity in Ifi205^K364ER299E^ (Fig.7C). Similar results were observed by EMSA using 20nM FAM labeled 60mer dsDNA (Fig.S5).

*Ifi205* or *Ifi205^K364ER299E^* was ectopically expressed in *Ifi205^-/-^ Mre11^ATLD1/ATLD1^* organoids followed by Western blot to assess IFN and chromatin markers. Ectopically expressing *Ifi205* resulted in much higher Stat1, macroH2A2 and Hp1-γ than that in organoids complemented with *Ifi205^K364ER299E^*, verifying that DNA binding by IFI205 underlies the ISG signature and chromatin changes in *Mre11^ATLD1/ATLD1^* MECs (Fig.7D).

### Mammary Tumorigenesis in Mre11^ATLD1/ATLD1^is Ifi205 Dependent

*Ifi205* deficiency reverts the oncogenic response of *Mre11^ATLD1/ATLD1^*to that of *WT* MECs, including the induction of heterochromatic marks and changes in chromatin accessibility. We reasoned that if these chromatin responses are tumor suppressive, then *Ifi205* deficiency in *Mre11^ATLD1/ATLD1^* MECs should reduce tumor incidence. To test this hypothesis*, WT, Mre11^ATLD1/ATLD1^* and *Ifi205^-/-^ Mre11^ATLD1/ATLD1^* organoids were implanted into cleared mammary fat pads of NOD-SCID mice. 6 weeks after implantation, 0.2mg/ml doxycycline water was fed to the mice to induce *neuT* expression. Successful implantation could be verified after 1 week of doxycycline feeding by luciferase imaging, whereas tumors were visible as large luciferase positive masses after oncogene activation (Fig.8A). Macroscopic images and H&E staining confirmed the existence of the primary tumor and lung metastasis from *Mre11^ATLD1/ATLD1^*organoid implantations (Fig.8B-C). 15 *WT,* 15 *Mre11^ATLD1/ATLD1^* and 17 *Ifi205^-/-^ Mre11^ATLD1/ATLD1^* organoid implanted mice were followed-up for 35 weeks after doxycycline induction. Tumor free survival in *Ifi205^-/-^ Mre11^ATLD1/ATLD1^* cohort after *neuT* activation was significantly longer than that in *Mre11^ATLD1/ATLD1^* cohort (Fig.8D, *Ifi205^-/-^ Mre11^ATLD1/ATLD1^* / *Mre11^ATLD1/ATLD1^*, HR=0.1496, p=0.0414; *WT* / *Mre11^ATLD1/ATLD1^*, HR=0.1146, p=0.0159; Log-rank test). To include early-stage tumors that are not palpable, mammary tissues harvested at the end the experiment were submitted for pathology analysis. Microscopic primary tumor incidence also decreased in *Ifi205^-/-^ Mre11^ATLD1/ATLD1^* implantations compared with *Mre11^ATLD1/ATLD1^* implantations (Fig.8E, *Mre11^ATLD1/ATLD1^*, 85.71%; *Ifi205^-/-^ Mre11^ATLD1/ATLD1^*, 23.53%, p=0.0010). These data show that ablation of *Ifi205* in *Mre11^ATLD1/ATLD1^* organoids suppresses oncogene-induced tumorigenesis in *Mre11^ATLD1/ATLD1^*organoid implantation.

**Figure 8.**
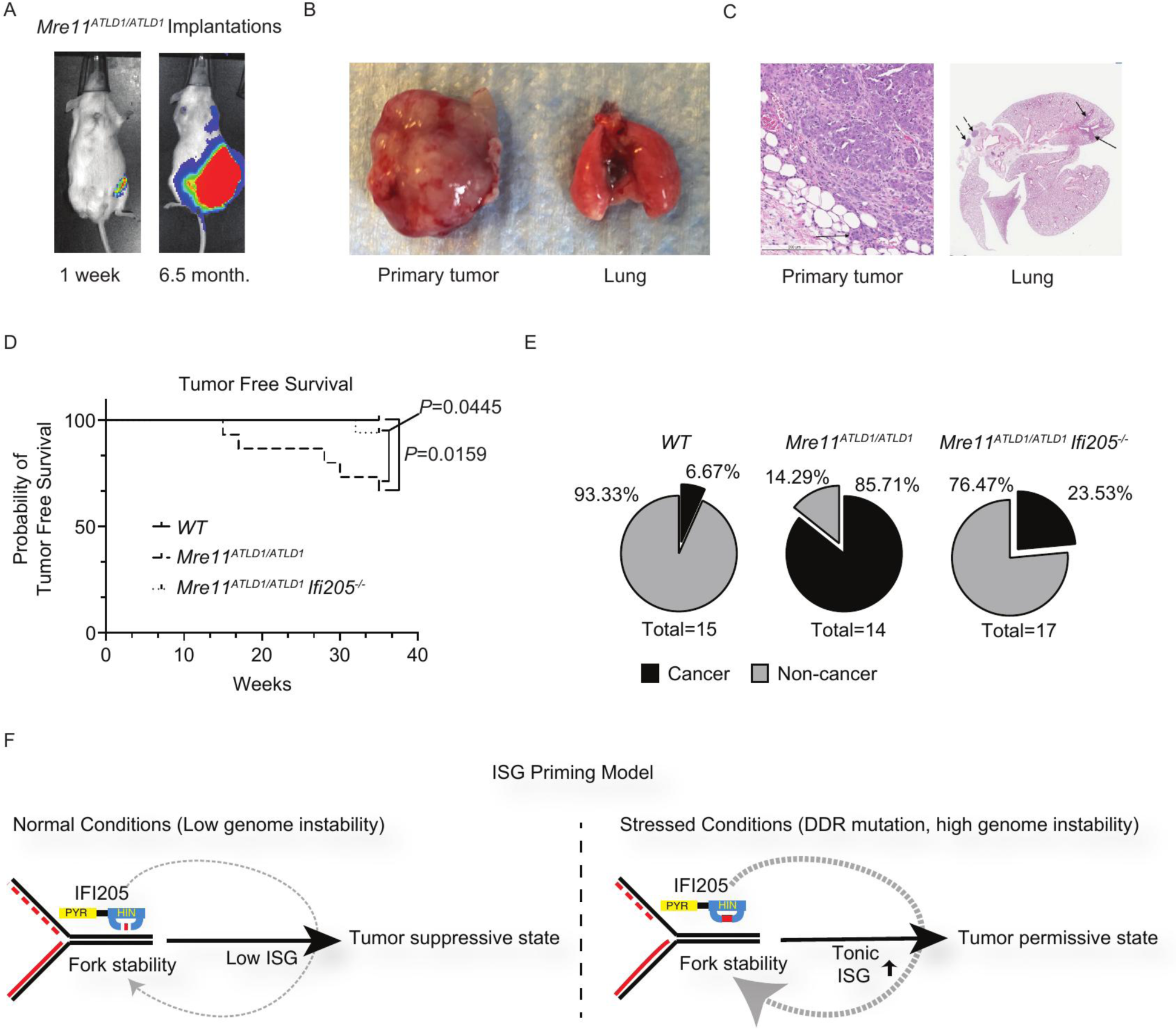
Knocking-out *lfi205* in *Mre11^ATLD1/ATLD1^* organoids suppresses oncogene-induced mammary tumorigenesis. (A) Luciferase images of *Mre11^ATLD1/ATLD1^* organoids implanted mouse at 1 week (left) and 6.5 months (right) after fed with 0.2mg/ml doxycycline water. (B) Macroscopic primary tumor (left) and lung sample (right) from *Mre11^ATLD1/ATLD1^* organoids implanted mouse at 7 months after fed with 0.2mg/ml doxycycline water. (C) H&E staining of samples from (B) (Arrows pointing to lung metastasis lesions). (D) Kaplan-Meier tumor free survival curves of *WT, Mre11^ATLD1/ATLD1^* and *lfi205^-/-^ Mre11^ATLD1/ATLD1^* organoids implanted mice after fed with 0.2mg/ml doxycycline water *(WT* n=15, *Mre11^ATLD1/ATLD1^* n=15, *lfi205^-/-^ Mre11^ATLD1/ATLD1^* n=17: lfi205^-/-^ *Mre11^ATLD1/ATLD1^ / Mre11^ATLD1/ATLD1^,* HR=0.1496, P=0.0445; *WT / Mre11^ATLD1/ATLD1^*, HR=0.1146, P=0.0159; Log-rank test). (E) Percentage of microscopic primary tumor incidence at the time of experiment termination (35 weeks after fed with 0.2mg/ml doxycycline water) in *WT, Mre11^ATLD1/ATLD1^* and *1/1205^-/-^ Mre11^ATLD1/ATLD1^* organoids implanted mice *(Mre11^ATLD1/ATLD1^* 85.71%, *lfi205^-/-^ Mre11^ATLD1/ATLD1^* 23.53%, P=0.0010, Fisher’s exact test). (F) Schematic model. DNA sensors such as IFI205 are poised to detect aberrant DNA structures at the DNA replication fork. Under normal conditions, such structures are infrequently encountered during unperturbed DNA replication, leading to low level of ISG transcription and tumor suppressive state. Under stressed conditions such as DDR mutation, constant engagement with aberrant DNA structures results in elevated ISG transcription and tumor permissive state, increasing susceptibility for oncogene-induced tumorigenesis. The ISG transcription program may also affect replication fork stability as a feedback loop.

## Discussion

In this study, we used primary mammary organoids to examine the mechanisms that underlie the suppression of oncogene induced breast cancer. Having previously established the importance of the Mre11 complex in suppressing *neuT* induced breast cancer via *in vivo* analyses^5^, the organoid system afforded several advantages over the *in vivo* system. It provides sufficient material for genomic, biochemical and chromatin analyses, and organoids could be manipulated *ex vivo* and subsequently reimplanted for *in vivo* development.

Activation of oncogene in organoids fully recapitulated previous *in vivo* observations, both with respect to the indices of DDR activation and to the deposition of heterochromatic marks. These outcomes were not seen in *Mre11^ATLD1/ATLD1^* organoids. Bulk RNA-Seq analyses were carried out prior to oncogene activation, 14, and 28 days after oncogene induction and revealed substantial differences between *WT* and *Mre11^ATLD1/ATLD1^* organoids; the trajectories of 5,644 genes differed between the two genotypes. Consistent with the fact that *WT* organoids exhibited indices of DDR activation and heterochromatin formation upon oncogene activation, transcription of DDR genes as well as chromatin remodeling and modifying genes was induced to a greater extent in *WT* than in *Mre11^ATLD1/ATLD1^* (Fig.S6). In addition, the transcriptional profiles of genes involved in lipid metabolism and extracellular matrix components were sharply down regulated by oncogene induction in *WT* organoids, whereas those genes were induced in *Mre11^ATLD1/ATLD1^* (Fig.2B). We speculate that this difference may partially account for the increased metastatic potential exhibited by *Mre11^ATLD1/ATLD1^* mammary tumors, but further analysis would be required to substantiate that interpretation.

As stated previously, the increased susceptibility of *Mre11^ATLD1/ATLD1^* could reflect the existence of a Mre11 complex dependent response to oncogene activation, or a state created by Mre11 complex hypomorphism in which the tumor suppressive response to oncogene induced biological changes is attenuated.

Certain aspects of the data presented herein support the former explanation. For example, in *WT* organoids, genes governing the DDR as well as chromatin modifiers are induced. In 53BP1 deficient mice, *neuT* activation did not lead to extensive hyperplasia or tumor susceptibility^5^. This argues that the DNA repair pathway(s) influenced by 53BP1 foci are not strongly tumor suppressive.

Indeed, the data more strongly support the latter idea—that Mre11 complex hypomorphism indirectly promotes *neuT* driven breast cancer, possibly via alteration of the chromatin landscape. The tonic ISG transcriptional signature observed at baseline in *Mre11^ATLD1/ATLD1^*organoids is associated with global changes in chromatin accessibility. Those changes are dependent on IFI205, inactivation of which largely restores the altered accessibility peaks in *Mre11^ATLD1/ATLD1^* to that of *WT* organoids (Fig.5H). IFI205 deficiency in *Mre11^ATLD1/ATLD1^* also reverts the response to *neuT* activation with respect to the induction of heterochromatic markers and changes in chromatin accessibility (Fig.6A and Fig.6B).

In contrast, IFI205 deficiency did not mitigate the DDR defects associated the *Mre11^ATLD1/ATLD1^*. Spontaneous chromosome aberrations in *Mre11^ATLD1/ATLD1^* and *Mre11^ATLD1/ATLD1^ Ifi205^-/-^* arose at similar frequencies, both of which were significantly higher than *WT* (Fig.S7). And the attenuation of γH2AX formation upon oncogene activation in *Mre11^ATLD1/ATLD1^* was not rescued by IFI205 deficiency (Fig.S8).

The data suggest that IFI205 dependent tonic ISG transcription in *Mre11^ATLD1/ATLD1^*mammary epithelium creates a permissive chromatin environment for oncogene-induced tumorigenesis. It also must be considered that the ISG transcriptional program involves myriad genes which either alone or in combination(s) could change other aspects of epithelial cell biology.

What is certain is that the effect is dependent on IFI205 DNA binding, which likely stems from the increased rate of spontaneous DNA damage ensuing from DNA replication in the context of Mre11 complex hypomorphism. Previous studies clearly indicate that the Mre11 complex is intimately associated with DNA replication, and that functional decrement of the complex causes DNA replication stress and chromatid breakage^46,51-54^. Moreover, IFI205 localizes to nascent DNA (likely DNA replication forks) in both unperturbed and stressed DNA replication conditions^46^; hence it is temporally and spatially situated to surveil the replication fork. Consistent with this idea, we found that gene amplification of *IFI16*, human orthologue of *Ifi205*, in breast cancer patients with high genome instability was associated with inferior overall survival using The Cancer Genome Atlas (TCGA) datasets (Fig.S9).

The relationship between chromatin status and ISG transcription is complicated^55-57^. One possibility is that ISG transcriptional activity is causative of the observed chromatin changes. In this scenario, chromatin remodelers may be regulated by ISG master transcription factors such as *Irf3* and *Irf7*. A second possibility is that chromatin changes arise first and cause the subsequent ISG transcription activation by increasing chromatin accessibility. The fact that *Irf7* and some chromatin regulators are differentially expressed in *Mre11^ATLD1/ATLD1^* organoids (Fig.3C and Fig.4A), without significant increase of chromatin accessibility in regions of IFN transcription factors or ISGs favors the former possibility (Full list of genes with increase chromatin accessibility in *Mre11^ATLD1/ATLD1^* organoids in Table S3).

IFN was first discovered as a major defense against viral infections^58^. ISGs that encode cytotoxic proteins are induced in the acute phase of signaling in response to genotoxic stress such as that induced by treatment with anthracyclines or ionizing radiation^59,60^. In contrast, chronic stimulation with low dose of IFN contributes to resistance to DNA damage and correlated with therapy resistance^61^. It is now acknowledged that the effects of IFN signaling on cancers are determined by the strength and duration of stimulation; whereas strong and acute IFN responses are cytotoxic, weak and chronic response promote cell survival^62^. In support of this conclusion, we found stronger IFN signature in human breast cancer tissues compared with matched normal tissues, which is potentially associated with the pro-tumor effects of ISG chronic activation as observed in *Mre11^ATLD1/ATLD1^*organoids (Fig.S10).

Recent findings have illuminated a role for innate immune signaling in maintaining genomic integrity through the combined activities of nuclear DNA sensing (IFI16 in humans is orthologous to IFI205 and IFI204) and the induction of ISG15, which we showed is required for replication for stability and for the recruitment of factors that promote replisome function^46,47,63^. On the basis of those observations and the data presented in this study, we propose a model wherein innate immune sensors and effectors surveil of the DNA replication fork under normal growth conditions. The model posits that DNA sensors such as HIN-200 family members are poised to detect aberrant DNA structures at the DNA replication fork that are likely to exist transiently but frequently during unperturbed DNA replication. Engagement of such structures leads to activation of an ISG transcriptional program that culminates in the modulation of chromatin states and replication fork stability. In effect, the model suggests that this circuitry, which evolved as an antiviral mechanism has been coopted to effect surveillance of the replication fork^64^ (Fig.8F).

In this regard, chronic activation of innate immune pathways may contribute to the cancer predisposition and clinical sequalae of many DDR-related mutations as a result of ISG transcriptional activity stimulated by aberrant DNA structures. Further characterization of chromatin changes associated with chronic ISG transcription may provide information regarding tumor initiation and suggest novel treatment modalities.

## Materials and methods

### Animal experiments

All animal studies were done in compliance with a protocol approved by the Institutional Animal Care and Use Committee of Memorial Sloan Kettering Cancer Center. *CAG-rtTA* transgenic mice and *TetO-NeuT* transgenic mice were kindly provide by Scott Lowe (Memorial Sloan Kettering Cancer Center, USA) and Lewis Chodosh (University of Pennsylvania), respectively^13,14^. *Mre11^ATLD1/ATLD1^* mice have been previously described^45^. Mice were raised in a pathogen-free facility and genotyped by PCR.

Cleared mammary fat pad organoid implantation was done in 3-4-week-old female NOD-SCID mice. Mammary fat pad clearance procedure was performed as previously described^65^. Briefly, the bridge between the 4^th^ and 5^th^ mammary glands was cut and the region of the mammary fat pad between the nipple and just after the proximal lymph node was removed. 1 million single cell / Matrigel mixture (1:1) from desired genotype was injected into the remaining portion of the epithelium-free mammary fat pad. The skin incisions were closed by would autoclips.

Luciferase imaging was performed six weeks after organoid implantation. Organoid implanted mice were fed with 0.2 mg/ml doxycycline water for one week before imaging. Image was taken using Ivis Optical Imaging System 10min after D-luciferin IP injection.

Mouse cohorts after successful implantation were palpated for the development of mammary tumors twice a week. Mice were euthanized using humane experimental endpoints or at the termination of experiments. At necropsy, mammary tissues and lung tissues were harvested and fixed in 4% paraformaldehyde. Samples were processed for paraffin embedding, H&E stanning and pathology analysis (HistoWiz).

### Primary mammary organoid culture

The 4^th^ mammary glands were harvested from 6-8-week-old female mice from desired genotype and incubated in Collagenase/Hyaluronidase medium (Stem Cell Technologies, #07912) and shaken at 37°C for 2h. The resulting digestion was spun down and washed twice with D10F medium (DMEM, 10%FBS). Cells were washed four more times with DMEM. Cells were resuspended in 3ml trypsin and incubated at 37°C for 10min. D10F was added to neutralize the trypsin. Cells was spun down and resuspended in 10U Dispase (Stem Cell Technologies, #07923) and 1000U DNase (Worthington) and incubated at 37°C for 10min. D10F was added and cells were passed through 40µm filter. Cells were spun down, embedded in Matrigel (Fisher Scientific, CB40230C) and overlaid with growth mammary organoid medium. For maintenance, organoids were culture in growth mammary organoid medium (basal mammary organoid medium plus 50ng/ml EGF (Peprotech, 315-09)) and passaged on a weekly basis using trituration with a fire polished glass Pasteur pipet or by trypsinization. Basal state organoids were cultured in basal mammary organoid medium for 1 day and oncogene activated organoids were cultured in doxycycline mammary organoid medium (basal mammary organoid medium plus 1μg/ml doxycycline (Millipore Sigma, D3072)) for indicated time. A full list of ingredients and concentrations is given in Table S4.

### Immunofluorescence

Organoids were washed twice with PBS and incubated with Cell Recovery Solution (Fisher Scientific, 354253) on ice for 1h. Organoids were fixed with 4% PFA at 4°C for 1 hour, then permeabilized using 1% triton-X100 in PBS for 1 hour at RT. Staining was performed in 0.3% Triton X-100 and 1% BSA in PBS overnight at 4°C on gently shaking. Antibodies used were listed in Table S5. Stained organoids were imaged using a SP8 confocal microscope.

### RNA extraction and quantitative PCR (qPCR)

Organoids were isolated from the Matrigel by multiple washes with ice cold PBS. Total RNA was extracted from the specified organoid using the RNeasy Mini Kit (Qiagen, 74104) with DNase treatment. cDNA was synthesized using RNA to cDNA EcoDry Premix (Takara, 639547). qPCR was performed using SsoAdvanced Universal SYBR Green Supermix (Biorad, 1725272) on a CFX384 real time system (BioRad). Target gene expression was normalized to *Gapdh* and relative expression determined with comparative CT method. Primers used are listed in Table S6.

### Whole cell extract western blot

Organoids were isolated from the Matrigel by multiple washes with ice cold PBS. Cells were lysed in RIPA buffer containing protease inhibitors (Sigma, 11873580001) and phosphatase inhibitors (Thermo Fisher Scientific, 78420) on ice and sonicated ten times for 30s at 30s intervals using a Bioruptor. Protein concentrations were quantified using a bicinchoninic acid (BCA) assay (Thermo Fisher, 23227). Lysates were denatured using 5X protein loading dye (300 mM Tris-HCl pH 6.8, 10% SDS, 0.5% bromophenol blue, 50% glycerol, 500 mM 2-mercaptoethanol). 20-30 μg of extract was loaded into gradient SDS-PAGE gels (Bio-Rad) and transferred to nitrocellulose membranes (Amersham Proton 0.2 μM NC #10600006). The membranes were blocked in 5% skimmed milk PBST and probed with the antibodies listed in Table S5. Membranes were developed using either ECL or ECL Prime (Amersham RPN2106/RPN2232).

### *Ifi205* knockout

*Ifi205* knockout organoids was generated using CRISPR-Cas9-mediated genome editing as described previously^66^. Briefly, *Ifi205* guide sequence (TGAAGCCGAAGATGAGACCT) was cloned into *PX458* (Addgene, 48138) using *Bbs*I restriction sites. The plasmid was nucleofected (Nucleofector II, Amaxa) into organoids and sorted for GFP positive cells 72h later. A second round of nucleofection was performed and sorted for GFP positive cells again. Cells were genotyped via sequencing and ICE analysis (www.synthego.com).

### Metaphase spreads

Organoids were treated with 100 ng/ml of KaryoMAX colcemid (Life technologies, 15212012) for 24 h and harvested. Organoids were washed with PBS to remove Matrigel and made into single cell suspension by trypsinization. Cells were swelled in 0.075 M KCl for 15 min at 37 °C and fixed in ice-cold 3:1(v/v) methanol: acetic acid at −20 °C overnight. Samples were dropped on slides and stained with 5% Giemsa (Sigma) at room temperature for 5 min. Following 3× washes in dH_2_O slides were dried for several hours at RT and mounted with Permount medium (Fisher Scientific). Metaphases were imaged on Olympus IX50 microscope with an Infinity 3 camera (Lumenera) using a 100X Objective.

### Recombinant protein expression and purification

Mutagenesis for *Ifi205^K364ER299E^* was performed in *pCMV6-Ifi205* (Origene plasmid, MR206355) using QuikChange Lightning Multi Site-Directed Mutagenesis Kit (Agilent, 200515). Primers used for mutagenesis are listed in Table S6. Sequences encoding *Ifi205* ORF and *Ifi205^K364ER299E^* ORF were amplified by PCR then subcloned between the BamHI and EcoRI restriction sites of *pGEX-6P-1*(Addgene). Ifi205 proteins were expressed as GST-Ifi205-FLAG-fusions in the *E. coli* strain BL21(DE). Proteins were purified by Glutathione Sepharose (GE Healthcare, GE17-0756-01) and cleaved by HRV-3C protease (Millipore Sigma, SAE0045) to remove GST tag. The resulting samples were injected into Superdex 200 columns (GE Healthcare) for size exclusion chromatography. Ifi205 proteins corresponding fractions were combined and purified by ANTI-FLAG M2 Affinity Gel (Sigma, A2220). A small portion of the final proteins were run through the Superdex 200 columns (GE Healthcare) again for quality check.

### Binding Assay

Oligonucleotides were refolded in 50mM Tris pH 7.5, 150mM NaCl and 1mM EDTA with slow cooling starting at 95 °C down to room temperature. The change in fluorescence anisotropy of the 60mer 5’ FAM-labeled oligonucleotides was measured to determine the relative binding affinities of Ifi205 proteins. The labeled oligonucleotides (50nM) were mixed in 10μl reaction volumes with the indicated Ifi205 proteins at concentrations ranging from 0 to 750nM in a reaction buffer containing 20 mM Tris, pH 7.5, 50 mM KCl, 0.5 mM TCEP, 10% glycerol, and 0.1% IGEPAL in a 384-well microplate. The binding data were collected on a SpectraMax M5 (Molecular Devices) using a 495 nm excitation wavelength and a 525 nm emission wavelength. Apparent K_D_ values were calculated from triplicate experiments using a model for receptor depletion and plotted using Prism 7 GraphPad Software. The model used was: *Y* = Af+ (Ab − Af)*((*L* + Kd+ *X*) − sqrt((sqr (− *L* − Kd − *X*)) − 4\**L*\**X*))/(2\**L*). Where *Y* is the anisotropy measured; *A*_b_ is the anisotropy at saturation (100%), *A*_f_ is the anisotropy of the free oligonucleotide, *L* is the fixed concentration of the oligonucleotide and *X* is the protein concentration.

### Electrophoretic mobility shift assay (EMSA)

Oligonucleotides were refolded in 50mM Tris pH 7.5, 150mM NaCl and 1mM EDTA with slow cooling starting at 95 °C down to room temperature. Binding reactions were performed in buffer containing 20 mM Tris, pH 7.5, 50 mM KCl, 0.5 mM TCEP, 10% glycerol, and 0.1% IGEPAL. Ifi205 proteins and 60mer 5’ FAM-labeled oligonucleotide at the indicated concentrations were incubated at room temperature for 30 min then loaded onto a 4% polyacrylamide non-denaturing gel. The gels were imaged Typhoon FLA9500 instrument (GE).

### Ifi205 proteins expression in organoids

Sequences encoding *Ifi205* ORF and *Ifi205^K364ER299E^*ORF were amplified by PCR then subcloned between the XbaI and BamHI restriction sites of *pHFUW-IRES-EGFP*. Lentivirus expressing *pHFUW-IRES-EGFP* were produced in HEK293T cells using packaging vectors *psPAX2* and *pVSV-*G. Viral supernatant was concentrated using PEG-it Virus Precipitation Solution (SBI, LV810A-1). Spinoculation was done at 600g for 1h at 32°C. Organoid-virus mixture was incubated for additional 6h at 37°C in a culture incubator and refreshed with growth mammary organoid medium. 2 weeks after infection, organoids with different Ifi205 proteins were sorted for similar intensity of GFP. *cGAMP quantification*

Organoids were washed with PBS to remove Matrigel and made into single cell suspension by trypsinization. Cells were thoroughly resuspended in 120 μL lysis buffer (20 mM Tris-HCl pH 7.7, 100 mM NaCl, 10 mM NaF, 20 mM β-glycerophosphate, 5 mM MgCl_2_, 0.1% Triton X-100, 5% glycerol) and lysed with a 28½ gauge needle. Lysates were incubated on ice for 30 min, centrifuged at 16,000g, 4° C for 10 min and cGAMP levels were quantified using the 2′3′cGAMP ELISA Kit (Arbor Assays, NC1595685) according to the manufacturer’s instructions.

### Quantification of cytoplasmic dsDNA

Organoids were washed with PBS to remove Matrigel and made into single cell suspension by trypsinization. Cytoplasmic extracts from live were isolated using NE-PER Nuclear and Cytoplasmic Extraction Reagents (ThermoFisher Scientific, 78833). dsDNA was quantified in cytoplasmic fraction using the SpectraMax Quant AccuClear Nano dsDNA Assay kit (Molecular devices, Part # R8357). The samples were read using SpectraMax M5 (Molecular Devices) plater reader.

### Bulk RNA-Seq analysis

Organoids were washed with PBS to remove Matrigel and pelleted cells were frozen until process started. Frozen cells were lysed in 1 mL TRIzol Reagent (ThermoFisher catalog # 15596018) and phase separation was induced with 200 µL chloroform. RNA was extracted from 350 µL of the aqueous phase using the miRNeasy Mini Kit (Qiagen catalog # 217004) on the QIAcube Connect (Qiagen) according to the manufacturer’s protocol. Samples were eluted in 34 µL RNase-free water. After RiboGreen quantification and quality control by Agilent BioAnalyzer, 500 ng of total RNA with RIN values of 7.2-10 underwent polyA selection and TruSeq library preparation according to instructions provided by Illumina (TruSeq Stranded mRNA LT Kit, catalog # RS-122- 2102), with 8 cycles of PCR. Samples were barcoded and run on a HiSeq 4000 in a PE50 run, using the HiSeq 3000/4000 SBS Kit (Illumina). An average of 61 million paired reads were generated per sample and the percent of mRNA bases averaged 79%. 50bp paired-end reads were aligned to the mm10 mouse reference genome using STATaligner. Quantification of genes annoted in Gencode vM2 was performed using FeatureCounts. Batch effects were corrected using RUVSeq if samples were submitted at multiple times. Differential expression analysis was done using the DESeq2 package. Differentially expressed genes were defined if they satisfied an adjusted-p < 0.01 and if the magnitude of foldchange >1.5. Canonical pathway analysis was performed by QIAGEN Ingenuity Pathway Analysis (QIAGEN IPA). GSEA was performed by fgsea package in R. GO analysis and KEGG pathway analysis were performed by GOstats package in R.

### Bulk ATAC-Seq analysis

Organoids were washed with PBS to remove Matrigel and made into single cell suspension by trypsinization. 50,000 freshly sorted live cells (DAPI negative) from each sample were used for ATAC-Seq as previously described^67^. Briefly, fresh cells were washed in cold PBS and lysed. The transposition reaction containing TDE1 Tagment DNA Enzyme (Illumina catalog # 20034198) was incubated at 37°C for 30 minutes. The DNA was cleaned with the MinElute PCR Purification Kit (Qiagen catalog # 28004) and material was amplified for 5 cycles using NEBNext High-Fidelity 2X PCR Master Mix (New England Biolabs catalog # M0541L). After evaluation by real-time PCR, 8-14 additional PCR cycles were done. The final product was cleaned by aMPure XP beads (Beckman Coulter catalog # A63882) at a 1X ratio, and size selection was performed at a 0.5X ratio. Libraries were sequenced on a HiSeq 4000 or NovaSeq 6000 in a PE50 or PE100 run, using the HiSeq 3000/4000 SBS Kit or NovaSeq 6000 S2 or S4 Reagent Kit (200 Cycles) (Illumina). An average of 64 million paired reads were generated per sample. ATAC-Seq data processing was performed as previously described^68^. In brief, paired-end reads were aligned to the mm10 genome using Bowtie2. Mapped fragments were shifted for Tn5 tagmentation and used for peak calling with MACS2. Peaks that were not reproducible (IDR < 0.01) and overlapping the ENCODE mm10 functional genomics regions blacklist were discarded to improve quality. Reproducible peaks from each sample were combined to create a genome-wide atlas of accessible chromatin regions. Reads aligned to the atlas peak regions were counted using Bedtools. Batch effects were corrected using RUVSeq if samples were submitted at multiple times. Differential accessibility of the peaks was assessed by DESeq2 to the count table. Peaks were defined as differentially accessible if they satisfied an adjusted-p < 0.05 and if the magnitude of foldchange >1.5. The heatmaps of differentially accessibly peaks were generated using Deeptools.

### Single-cell RNA-Seq analysis

Organoids were washed with PBS to remove Matrigel and made into single cell suspension by trypsinization. Cells were incubated with Fc receptor blocker (BD, 553142) and multiplexed with TotalSeq-A (BioLegend). Freshly sorted live single cells (DAPI negative) from each sample were used for single-cell RNA-Seq on Chromium instrument (10X genomics) following the user guide manual for 3′ v3.1. In brief, FACS-sorted cells were washed once with PBS containing 1% bovine serum albumin (BSA) and resuspended in PBS containing 1% BSA to a final concentration of 700–1,300 cells per μl. The viability of cells was above 80%, as confirmed with 0.2% (w/v) Trypan Blue staining (Countess II). Cells were captured in droplets. Following reverse transcription and cell barcoding in droplets, emulsions were broken and cDNA purified using Dynabeads MyOne SILANE followed by PCR amplification per manual instruction. Between 10,000 to 20,000 cells were targeted for each sample. Samples were multiplexed together on one lane of 10X Chromium (using Hash Tag Oligonucleotides - HTO) following previously published protocol^69^. Final libraries were sequenced on Illumina NovaSeq S4 platform (R1 – 28 cycles, i7 – 8 cycles, R2 – 90 cycles). The cell-gene count matrix was constructed using the Sequence Quality Control (SEQC) package^69^. Viable cells were identified on the basis of library size and complexity, whereas cells with >20% of transcripts derived from mitochondria were excluded from further analysis. UMAP and PHATE were performed using Scanpy in Python. Lineage signature scores were generated using signature genes previously described^17^. ISG signature score were generated using genes from GO Terms (response to interferon-alpha, response to interferon beta, response to interferon gamma). Chromatin modifiers and remodelers used in PHATE were listed in Table S7.

### TCGA data analysis

Mutation counts, *IFI16* amplification status and survival metrics related to Breast Invasive Carcinoma (TCGA, PanCancer Atlas) were retrieved from cBioPortal (https://www.cbioportal.org). Patients with mutation counts exceeding either 130 or 140 were categorized into subgroups based on the absence and presence of *IFI16* amplification. Survival analysis and log-rank tests were carried out with the survival package in the R environment.

Transcriptome profiles and clinical data related to breast invasive carcinoma were retrieved from TCGA using TCGAbiolinks package in R. DESeq2 was used to generate IFN signature score (Human Gene Set: REACTOME_INTERFERON_SIGNALING) in primary tumor samples and matched normal samples. Paired t-test was performed to compare the difference between the two groups.

## Acknowledgements

We are grateful to all the members of the Petrini laboratory for helpful discussions. We thank Tom Kelly and Andy Koff for critical reading of the manuscript. We thank Wouter Karthaus, Wytse Bruinsma, Francisco Barriga and Francisco Sanchez-Rivera for valuable advice and technical assistance. We acknowledge Ronan Chaligné, the members of MSKCC Single Cell Analytics Innovation Laboratory and the members of MSKCC Integrated Genomics Operation Core for RNA sequencing, ATAC sequencing and analysis. This work was supported by NIH GM56888, GM59413, R35GM136278 (J.H.J.P) and the MSK Cancer Center Core Grant P30 CA008748.

---

**Figure S1.**
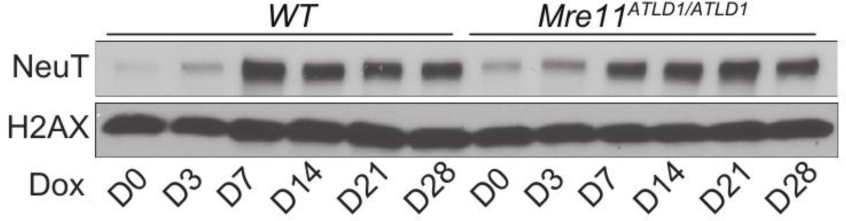
*neuT* induction in *WT* and *Mre11^ATLD1/ATLD1^* organoids. Western blot of whole cell extracts taken from H/T and *Mre11^ATLD1/ATLD1^* organoids at indicated timepoints after adding 1μg/ml doxycycline.

**Figure S2.**
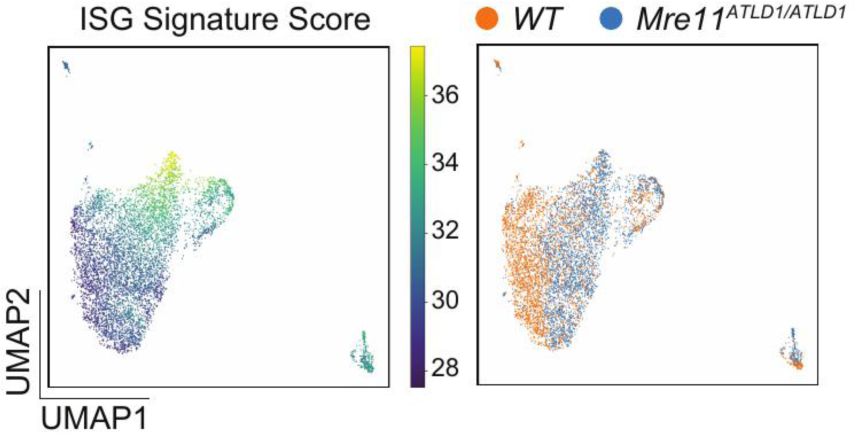
Up-regulation of ISG signature is a common feature in *Mre11^ATLD1/ATLD1^* organoids. UMAP plot color-coded by ISG signature score (left) and genotype (right) from single cell RNA-Seq in *WT* and *Mre11^ATLD1/ATLD1^* mammary organoids.

**Figure S3.**
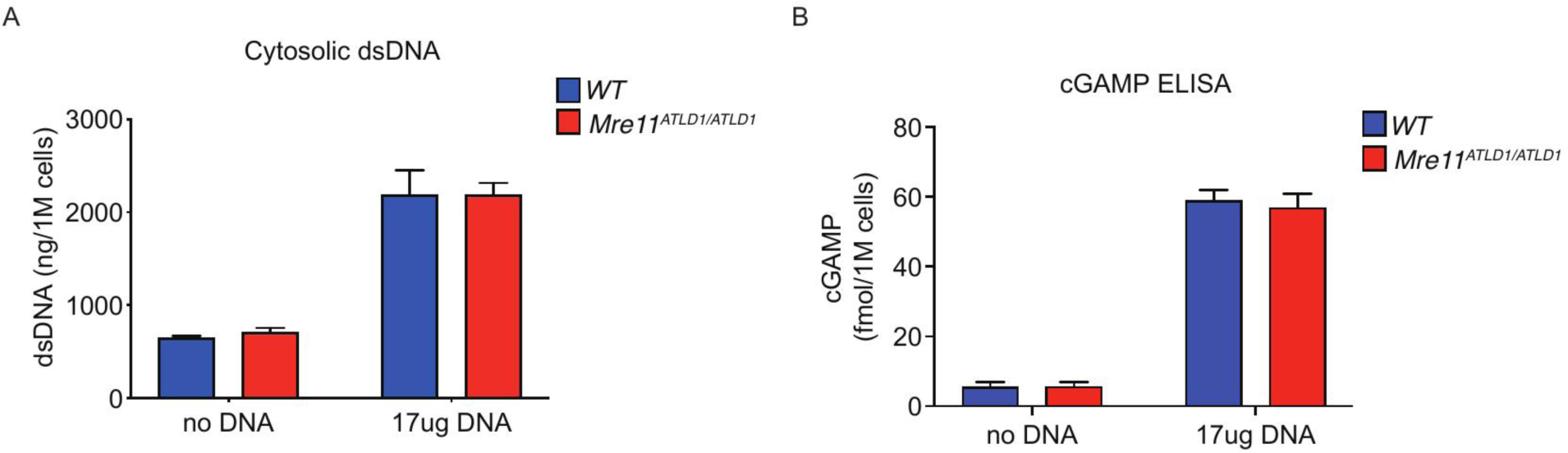
eGas-Sting DNA sensing pathway is not more active in *Mre11^ATLD1/ATLD1^* organoids compared with *WT* organoids. (A) Measurement of cytoplasmic dsDNA concentration in *WT* and *Mre11^ATLD1/ATLD1^* organoids (left), and in WT and *Mre11^ATLD1/ATLD1^* organoids after transfection of 17ug plasmid DNA as positive controls (right) *(WT* 643.0ng/1M cells, *Mre11^ATLD1/ATLD1^* 702.2ng/1M cells, t-test, p = 0.0130). (B) Measurement of Cyclic guanosine monophosphate-adenosine monophosphate (cGAMP) concentration by ELISA in *WT* and *Mre11^ATLD1/ATLD1^* organoids (left), and in *WT* and *Mre11^ATLD1/ATLD1^*organoids after transfection of Vug plasmid DNA as positive controls (right) *(WT* 5.383 fmol/1M cells, *Mre11^ATLD1/ATLD1^* 5.633 fmol/1M cells, t-test, p = 0.6336).

**Figure S4.**
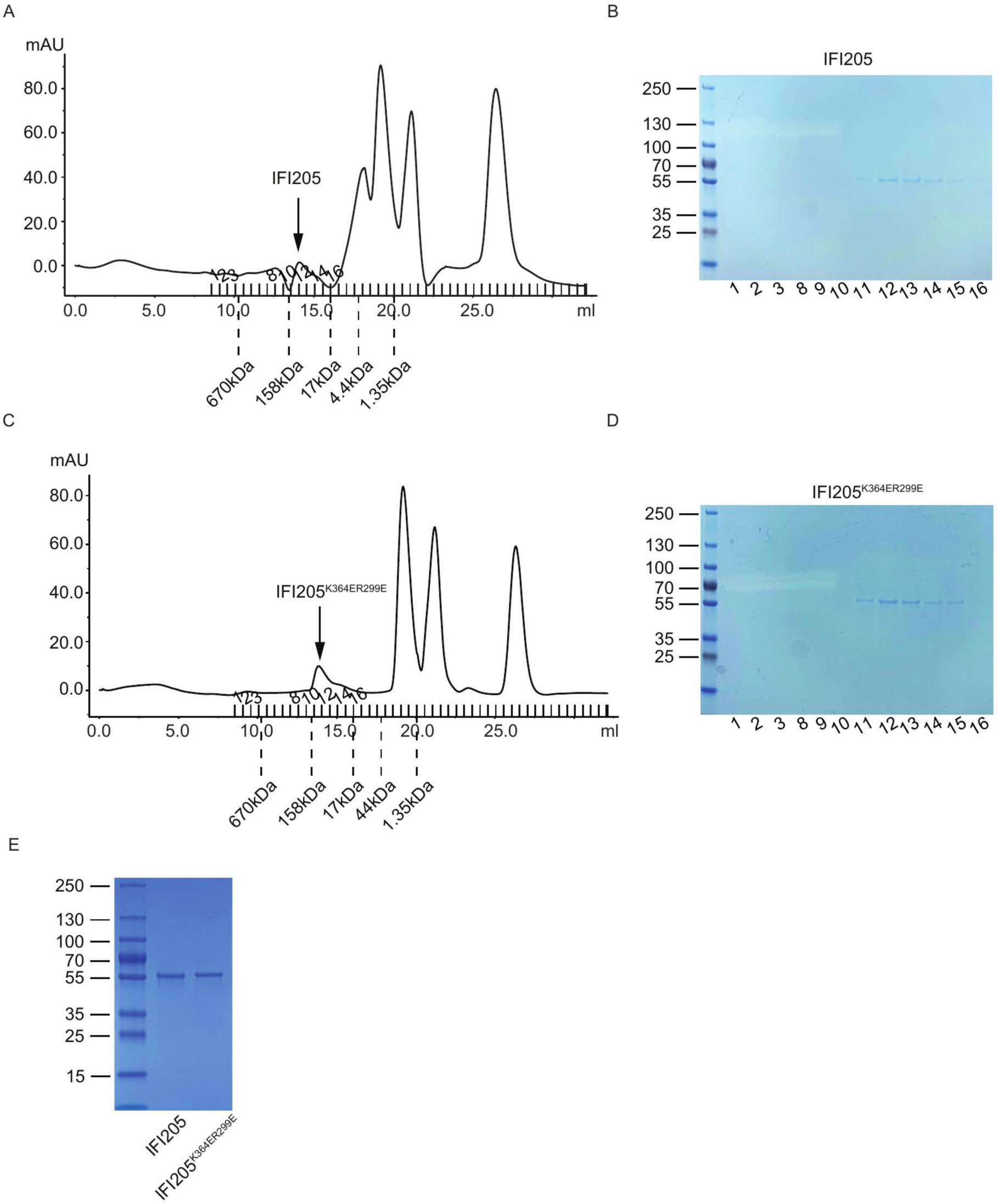
Recombinant IFI205 and IFI205^K364ER299E^ proteins were purified. (A) Size exclusion chromatography (SEC) of purifed recombinant IFI205 protein. Molecular weights were indicated acording to standard samples. (B) Commassie staining after SDS-PAGE using fractions from SEC. Fraction 1-3, 8-16 from (A) were loaded. (C) SEC of purifed recombinant ipi205^K364ER299E^ protein. Molecular weights were indicated acording to standard samples. (D) Commassie staining after SDS-PAGE using fractions from SEC. Fraction 1-3, 8-16 from (C) were loaded. (E) Commassie staining after SDS-PAGE for purified recombinant FLAG-tagged IFI205 and IFI205^K364ER299E^ proteins.

**Figure S5.**
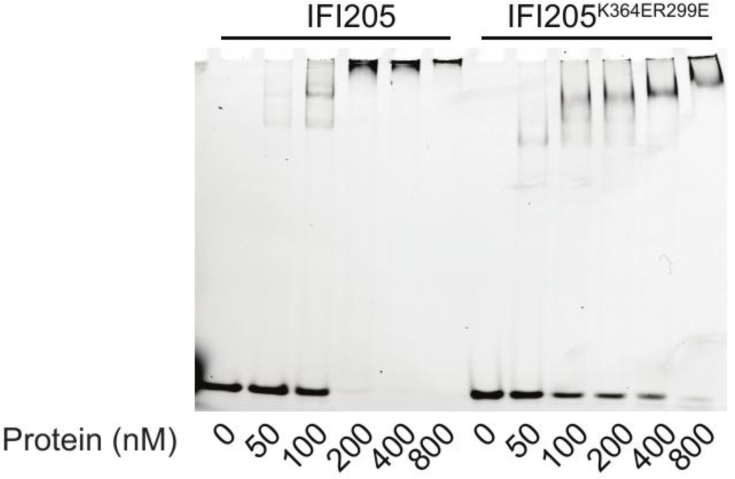
IFI205^K364ER299E^ shows deficient DNA binding compared with IFI205. Native electrophoretic mobility shift assay (EMSA) using 60mer FAM-labeled dsDNA at 20nM and increasing amount of recombinant IFI205 or IFI205^K364ER299E^ protein.

**Figure S6.**
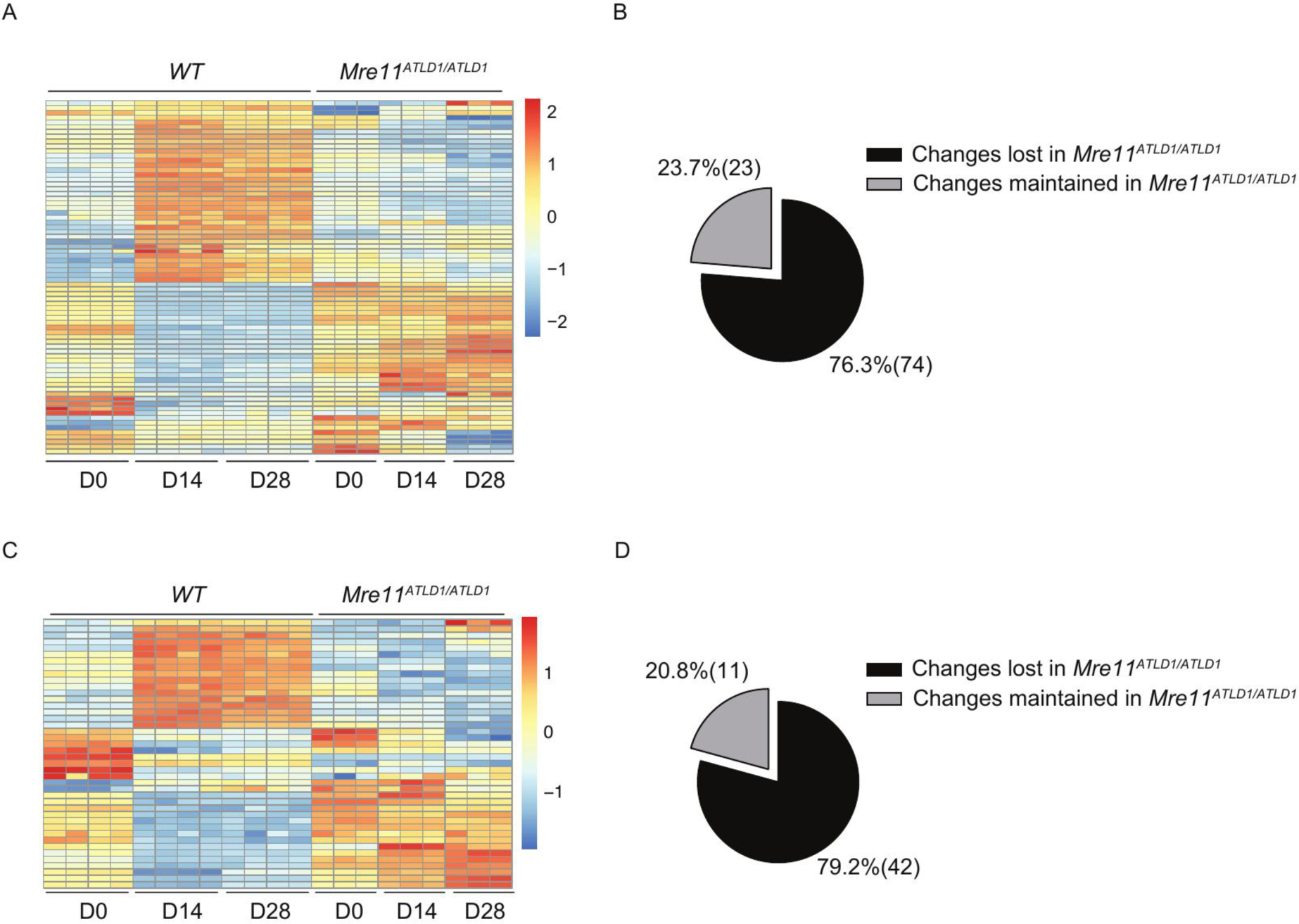
Majority of oncogene-induced changes of DDR genes and chromatin regulators observed in *WT* organoids are lost in *Mre11^ATLD1/ATLD1^* **organoids**. (A) Heatmap of DDR genes that were induced in *WT* organoids after oncogene activation but not in *Mre11^ATLD1/ATLD1^* organoids from RNA-Seq (padj ≤ 0.01 and |FC| ≥ 1.5). (B) Percentage of oncogene-induced DDR genes in *WT* organoids that are maintained or lost in *Mre11^ATLD1/ATLD1^* organoids from RNA-Seq (padj ≤ 0.01 and |FC| ≥ 1.5). (C) Heatmap of chromatin regulators that were induced in *WT* organoids after oncogene activation but not in *Mre11^ATLD1/ATLD1^* organoids from RNA-Seq (padj ≤ 0.01 and |FC| 2 1.5). (D) Percentage of oncogene-induced chromatin regulators in *WT* organoids that are maintained or lost in *Mre11^ATLD1/ATLD1^* organoids (padj ≤ 0.01 and |FC| ≥ 1.5).

**Figure S7.**
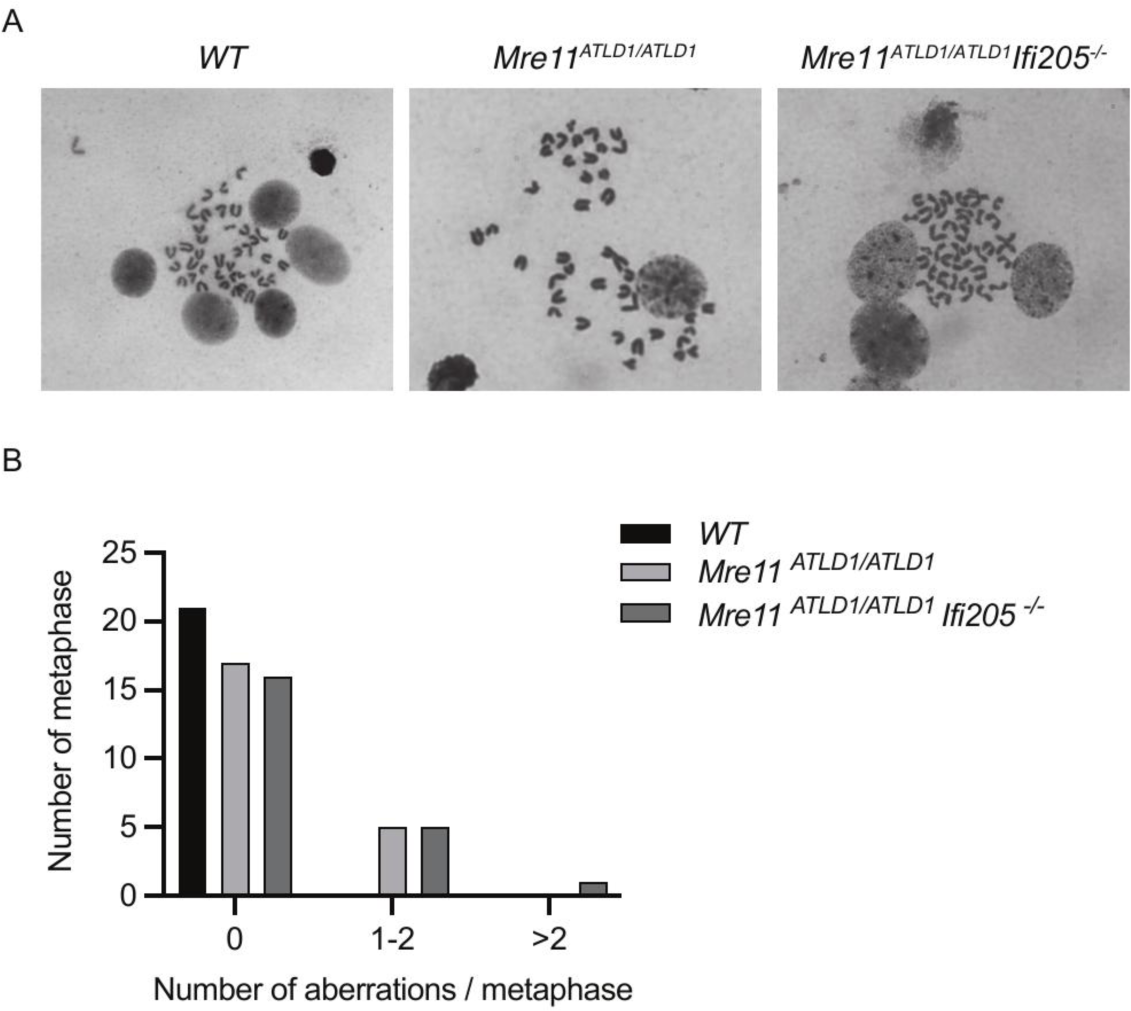
Inactivation of *lfi205* doesn’t rescue DNA damage in *Mre11^ATLD1/ATLD1^* organoids. (A) Example images of metaphase spreads in *WT, Mre11^ATLD1/ATLD1^* and *lfi205^-/-^Mre11^ATLD1/ATLD1^* organoids. (B) Quantification of metaphase spreads in A (21 *WT,* 22 *Mre11^ATLD1/ATLD1^*, 22 *lfi205^-/-^Mre11^ATLD1/ATLD1^, WT* vs *Mre11^ATLD1/ATLD1^*, p = 0.0201; *Mre11^ATLD1/ATLD1^* vs *lfi205^-/-^ Mre11^ATLD1/ATLD1^* p _=_ 0 5Q74; Chi-square test).

**Figure S8.**
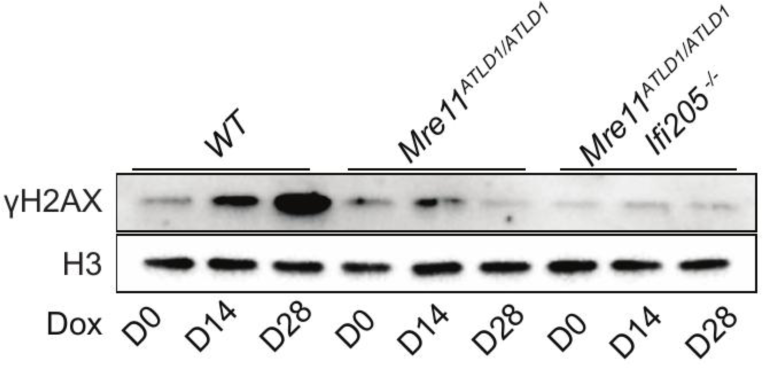
Inactivation of *lfi205* doesn’t rescue oncogene-induced DDR marker in *Mre11^ATLD1/ATLD1^* organoids. (A) Western blot of whole cell extracts taken from *WT* and *Mre11^ATLD1/ATLD1^*organoids at indicated timepoints after adding 1μg/ml doxycycline.

**Figure S9.**
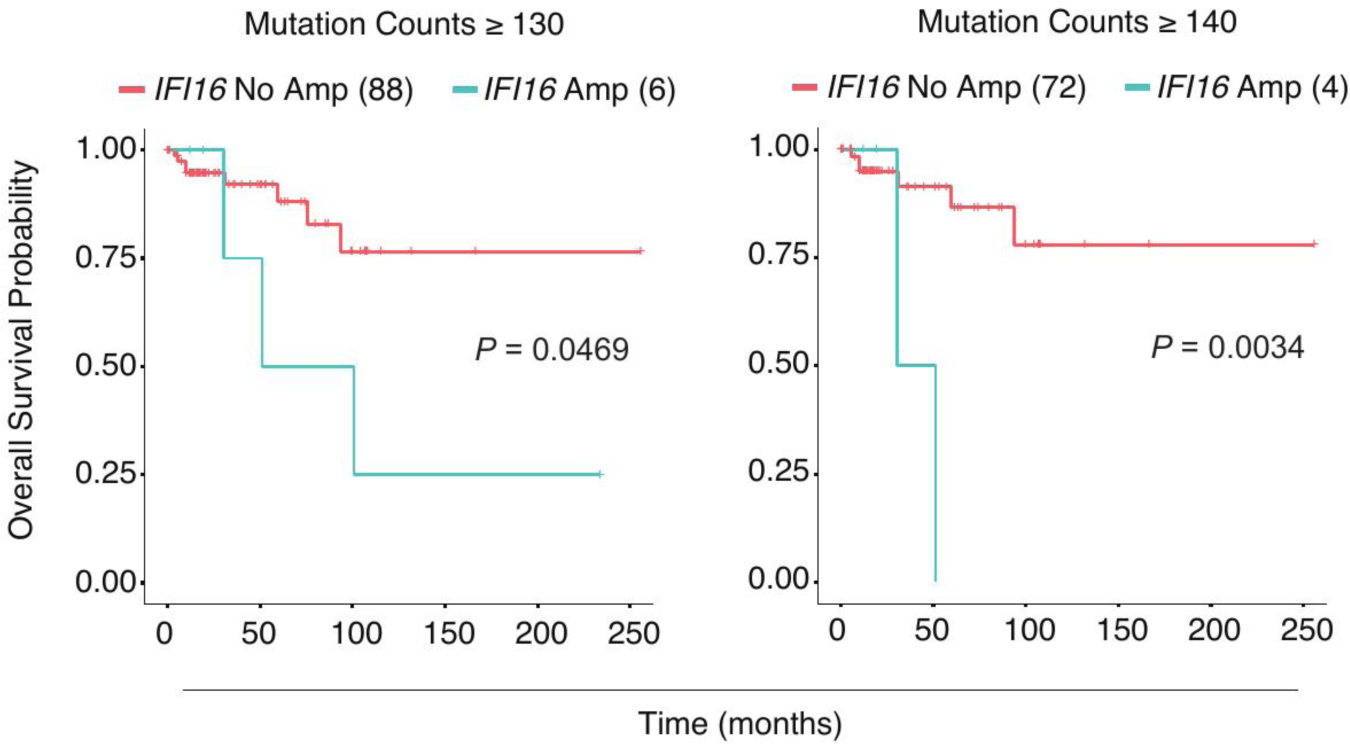
*IFI16* amplification shows inferior overall survival in breast cancer patients with high genome instability. (A) Overall survival curves for patients with more than 130 mutation counts in the presence or absence of *IFI16* amplification from Breast Invasive Carcinoma (TCGA, PanCancer Atlas) dataset (P = 0.0469, Log-rank test). (B) Overall survival curves for patients with more than 140 mutation counts in the presence or absence of *IFI16* amplification from Breast Invasive Carcinoma (TCGA, PanCancer Atlas) dataset. (P = 0.0034, Log-rank test).

**Figure S10.**
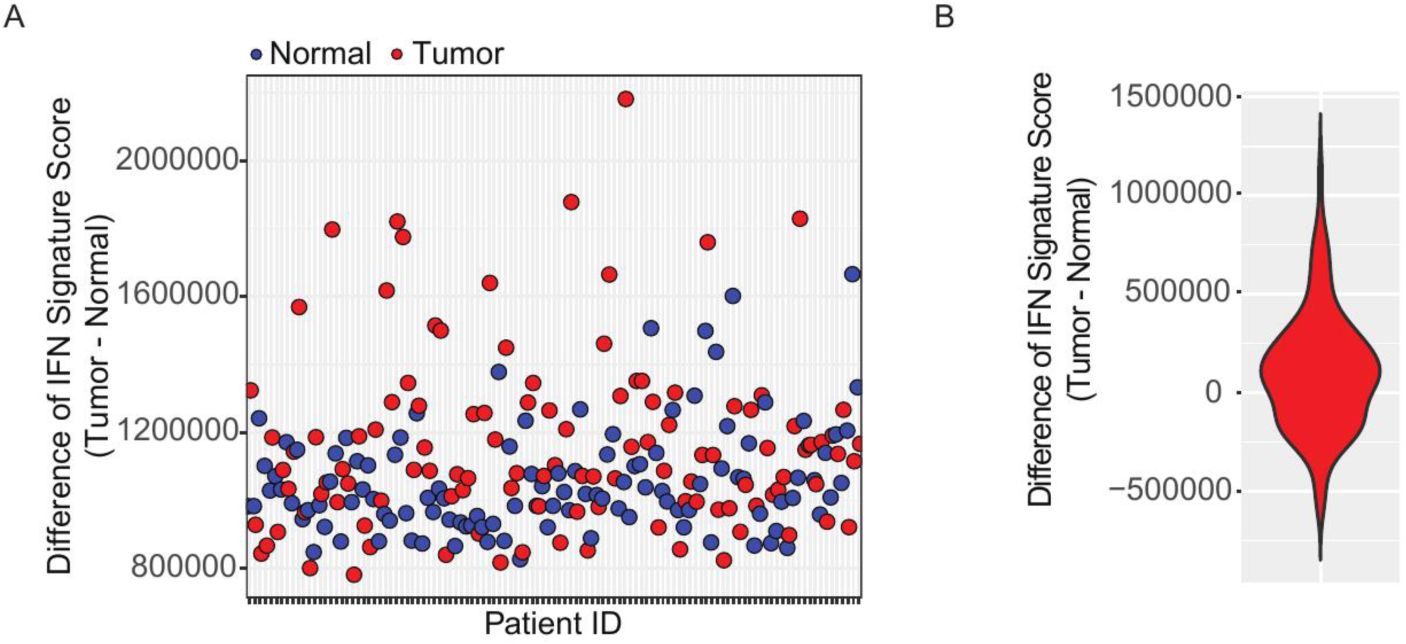
Human breast cancer tissues show elevated IFN signature score compared with matched normal breast tissues. (A) IFN signature score in 113 pairs of breast cancer tissues and their matched normal tissues from The Cancer Genome Atlas (TCGA) database. (B) Pairwise difference of IFN signature score in (A) (Tumor - normal, paired t-test, P=0.000368).

